# Activity-by-Contact model of enhancer specificity from thousands of CRISPR perturbations

**DOI:** 10.1101/529990

**Authors:** Charles P. Fulco, Joseph Nasser, Thouis R. Jones, Glen Munson, Drew T. Bergman, Vidya Subramanian, Sharon R. Grossman, Rockwell Anyoha, Tejal A. Patwardhan, Tung H. Nguyen, Michael Kane, Benjamin Doughty, Elizabeth M. Perez, Neva C. Durand, Elena K. Stamenova, Erez Lieberman Aiden, Eric S. Lander, Jesse M. Engreitz

## Abstract

Mammalian genomes harbor millions of noncoding elements called enhancers that quantitatively regulate gene expression, but it remains unclear which enhancers regulate which genes. Here we describe an experimental approach, based on CRISPR interference, RNA FISH, and flow cytometry (CRISPRi-FlowFISH), to perturb enhancers in the genome, and apply it to test >3,000 potential regulatory enhancer-gene connections across multiple genomic loci. A simple equation based on a mechanistic model for enhancer function performed remarkably well at predicting the complex patterns of regulatory connections we observe in our CRISPR dataset. This Activity-by-Contact (ABC) model involves multiplying measures of enhancer activity and enhancer-promoter 3D contacts, and can predict enhancer-gene connections in a given cell type based on chromatin state maps. Together, CRISPRi-FlowFISH and the ABC model provide a systematic approach to map and predict which enhancers regulate which genes, and will help to interpret the functions of the thousands of disease risk variants in the noncoding genome.

---

DNA elements in the human genome called enhancers control how different combinations of genes are expressed in different cell types and states, and harbor thousands of genetic variants that influence risk for common diseases. A major challenge in interpreting the functions of these variants is to map enhancer-gene connections: Which enhancers regulate which genes in which cell types, and with what quantitative effects?

Studies of individual enhancers and genes have shown that these connections can be complex: multiple enhancers can regulate a single gene, a single enhancer can regulate multiple genes across long genomic distances, and the network of enhancer-gene connections appears to rewire across cell types (*1, 2*). The mechanisms that give rise to this complexity remain poorly understood. One possibility (“biochemical specificity”) is that a given enhancer can regulate only the promoters that have complementary combinations of compatible transcription factors (TFs) (*3–6*). In a few cases it has been shown that specific TF-TF interactions are required for an enhancer to regulate a promoter (*7, 8*). Another possibility is that enhancer-gene interactions depend primarily on the 3D architecture of the genome, such as topological domains (*9, 10*) or chromatin loops (*1, 11, 12*). In a few cases it has been shown that manipulating enhancer-promoter contacts can affect gene expression (*13–15*). Various studies have integrated aspects of transcription factor binding and 3D architecture to attempt to predict enhancer-gene regulation (*16–19*). Yet, it has been difficult to evaluate these models or discover new ones because we have lacked efficient ways to study the regulatory effects of large numbers of enhancers in the genome.

We set out to map the effects of many putative enhancers on gene expression and thereby learn general rules to predict enhancer-gene connections across many cell types. We and others have recently developed high-throughput methods that use CRISPR to perturb noncoding elements in their native genomic locations to measure their effect on a target gene (*17, 20–24*). However, these methods have had two major limitations: (i) they cannot be readily applied to any target gene (they require that a gene has a phenotype that is well suited for multiplex screening, such as affecting cell proliferation, or is engineered to facilitate such screening, for example by introduction of a reporter construct under the control of its promoter in the genome) and (ii) they do not directly read out RNA levels. Related approaches using single cell RNA-Seq to quantify the effects of CRISPR perturbations (*25–29*) do not currently have sufficient sensitivity to detect small effects on most individual genes.

To overcome these limitations, we developed an approach called CRISPRi-FlowFISH to perturb hundreds of noncoding elements in parallel and quantify their effects on the expression of an RNA of interest (Fig. 1A; Fig. S1). In this approach, we design a library of guide RNAs (gRNAs) targeting a large collection of candidate regulatory elements, transduce the library into a population of cells expressing KRAB-dCas9 (on average 1 gRNA per cell), and induce KRAB-dCas9 expression for 48 hours. To measure the effects of candidate elements on the expression of a gene of interest, we: (i) use fluorescence in situ hybridization (FISH) to quantitatively label single cells according to their expression of an RNA of interest; (ii) sort labeled cells with fluorescence-activated cell sorting (FACS) into 6 bins based on RNA expression; (iii) use high-throughput sequencing to determine the abundance of each gRNA in each bin; (iv) and use this information to infer the effect of each gRNA on RNA expression. To assess quantitative effects and statistical significance, we calculate the average the effects of all gRNAs within each candidate element (Fig. S2A,B) and compare to hundreds of negative control gRNAs in the same screen.

**Fig. 1.**
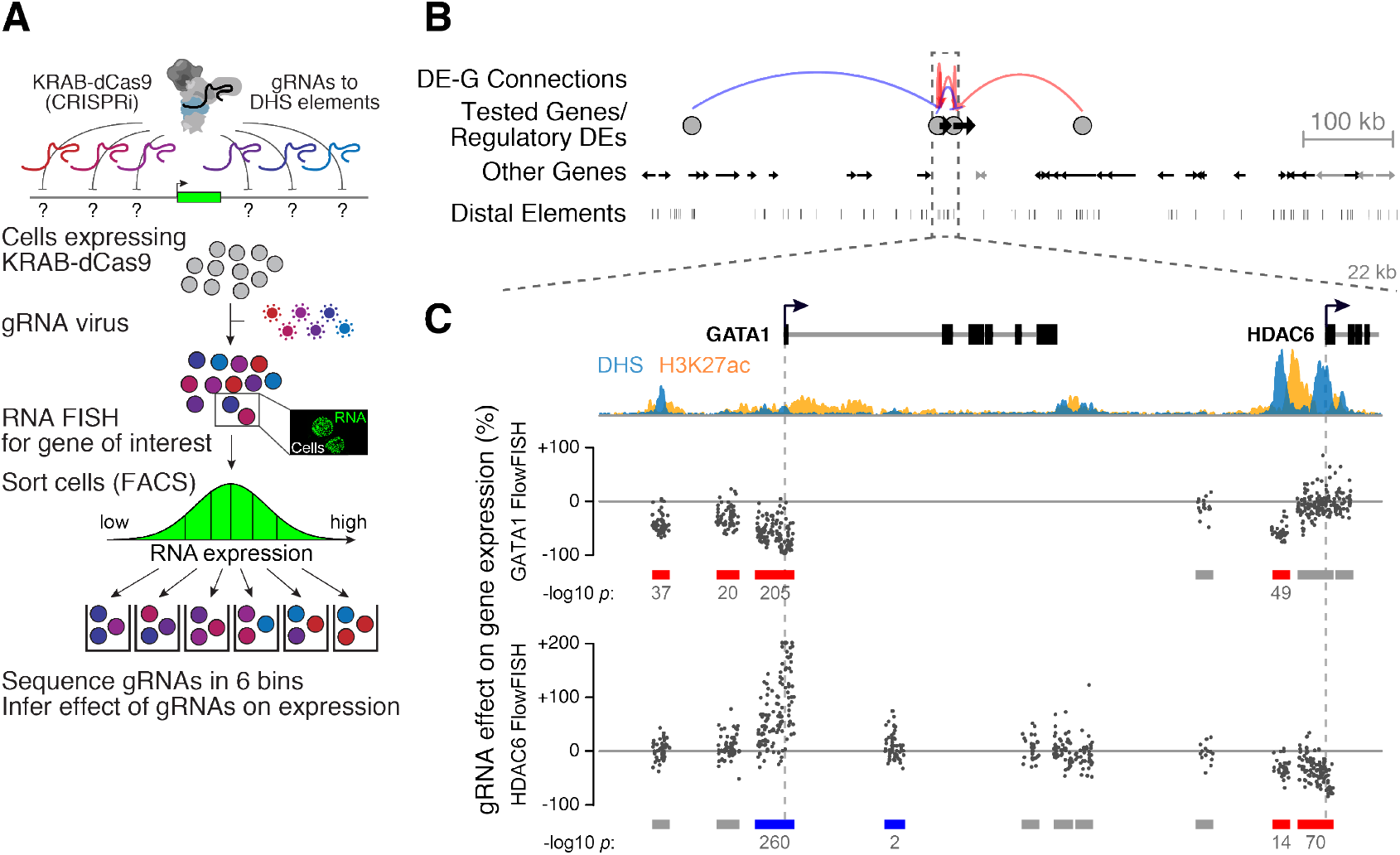
CRISPRi-FlowFISH identifies regulatory elements for *GATA1* and *HDAC6*. (**A**) CRISPRi-FlowFISH method for identifying gene regulatory elements. Cells expressing KRAB-dCas9 are infected with a pool of gRNAs targeting DHS elements near a gene of interest, labeled using RNA FISH against that gene, and sorted into bins of fluorescence signal by FACS. The quantitative effect of each gRNA on the expression of the gene is determined by sequencing the gRNAs within each bin. Inset: example of K562 cells labeled for *RPL13A*. (**B**) Distal elements affecting *GATA1* and *HDAC6* expression in K562 cells. Genes expressed in K562 cells are shown in black; those not expressed are shown in grey. Red arcs denote activation, blue arcs denote repression. See Fig. S3A for the full tested region spanning 4 Mb. (**C**) Close-up on region containing *GATA1* and *HDAC6*. Points represent the effect on gene expression of a single gRNA. *HDAC6* vertical axis capped at 200%. Grey, red, and blue bars: DHS elements in which CRISPRi leads to no detectable change (grey), a significant decrease (red) or increase (blue) in expression as measured by CRISPRi-FlowFISH. DHS elements in the gene body of the assayed gene are excluded from analyses because recruitment of KRAB-dCas9 to these sites may directly interfere with transcription.

To validate the approach, we first used CRISPRi-FlowFISH to identify elements that regulate the expression of *GATA1* in K562 human erythroleukemia cells. We performed replicate CRISPRi-FlowFISH screens using a probeset against *GATA1* and tested the functions of 127 candidate elements spanning 4 Mb (Fig. 1B,C; Fig. S3A). Replicate screens produced highly correlated estimates for the effect sizes of each element on *GATA1* expression (Pearson *R* = 0.95 for significant elements, Fig. S2C). As expected, these screens identified the three elements that we previously found to regulate *GATA1* (*17*), and we confirmed for individual gRNAs that the effects on gene expression estimated from CRISPRi-FlowFISH agreed with RT-qPCR measurements (Pearson *R* = 0.93, Fig. S1F). We note that these experiments do not distinguish between *cis* and *trans* effects (see Note S1).

To generate a large enhancer perturbation dataset, we used CRISPRi-FlowFISH in K562 cells to test a total of 3744 candidate regulatory element-gene pairs. Specifically, we designed FlowFISH assays for 28 genes in 5 genomic regions (spanning 1.1-4.0 Mb) and CRISPRi gRNAs against all DNase hypersensitive (DHS) elements in K562s within 450 kb of any of the genes (108 to 202 elements per gene for a total of 742 unique elements). The 28 genes included some with erythroid lineage-specific expression (*e.g., GATA1*) and some that are ubiquitously expressed (*e.g., RAB7A*), and were selected (after testing FlowFISH probesets for 51 genes) as those genes with probesets that met stringent criteria for both specificity and statistical power (Fig. S4, see Methods). We had >80% power to detect an effect on gene expression of 25% for all 28 genes and as low as 10% effects for 3 genes (Fig. S4C, see Methods). We analyzed these CRISPRi-FlowFISH data together with data from an additional 380 candidate regulatory element-gene pairs from previous CRISPR-based experiments in K562 cells, including our previous CRISPRi tiling proliferation screen in the *MYC* locus (*17, 23, 29–35*).

In total, our dataset included 3010 candidate distal element-gene (DE-G) pairs (where the targeted element is located >500 bp from a TSS) and 1114 distal promoter-gene (DP-G) pairs (where the targeted element is located <500 bp from a TSS). Here we focused on DE-G pairs, and analyzed DP-G pairs separately because we and others have found promoters can affect the expression of nearby genes through a variety of mechanisms beyond that of cis-acting enhancers (Note S2).

These perturbation-based maps uncovered complex connections wherein individual enhancers regulated up to 5 genes, individual genes were regulated by up to 12 distal elements, and in some cases enhancers appeared to “skip” over proximal genes to regulate more distant ones (Fig. 2; Fig. S3; Fig. S5). Of the 3010 DE-G pairs tested, 122 involved a significant effect on gene expression at a false discovery rate (FDR) < 0.05. The effect was activating in 80% of cases and repressing in 20% of cases (98 vs. 24), with absolute effect sizes ranging from 5%-93% (median: 24%; Table S3). Of 818 distinct DEs studied, 79 (10%) detectably regulated at least one gene in our dataset.

**Fig. 2.**
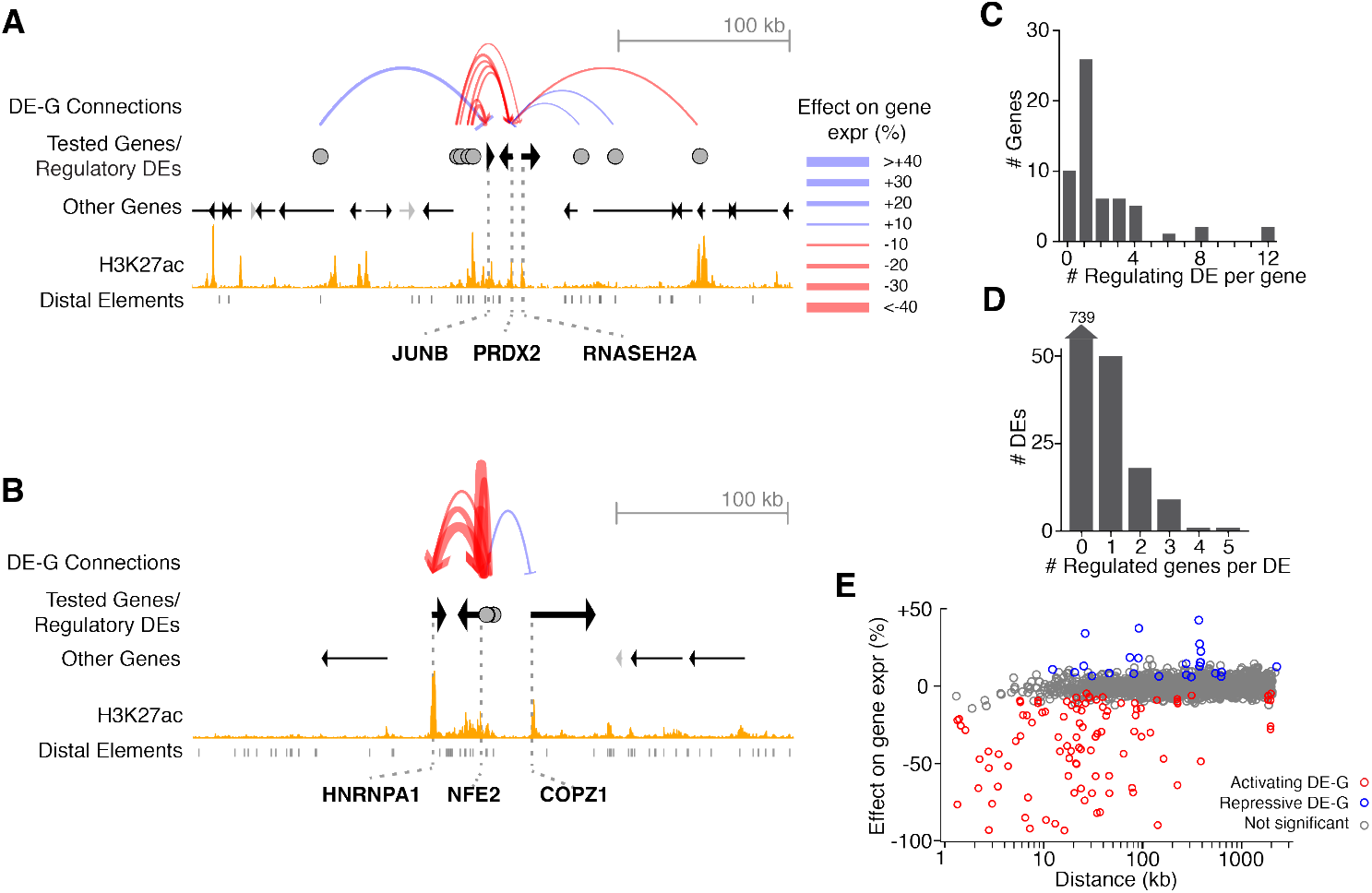
CRISPRi-FlowFISH produces regulatory maps of DE-G connections in multiple loci. (**A**) Example of CRISPRi-FlowFISH screen data. DE-G connections are elements affecting the expression of *JUNB, PRDX2*, and *RNASEH2A* in CRISPRi-FlowFISH screens in K562 cells. Red arcs denote activation, blue arcs denote repression. The width of the arc corresponds to the effect size. Distal elements are DHS peaks. Tested genes refer to genes for which we performed CRISPRi-FlowFISH experiments. See Fig. S3B for the full tested region spanning 1.4 Mb. (**B**) Same as (A) for the genes *HNRNPA1, NFE2*, and *COPZ1*. See Fig. S3C for the full tested region spanning 1.2 Mb. (**C**) Histogram of the number of distal elements affecting each gene in our dataset. (**D**) Histogram of the number of genes affected by each distal element tested in our dataset. (**E**) Comparison of genomic distance with observed changes in gene expression upon CRISPR perturbations. Each dot represents one tested DE-G. Red dots: connections where perturbation resulted in a decrease in the expression of the tested gene. Blue dots: perturbation resulted in an increase. Grey dots: had no significant effect. Panels (C-E) include both FlowFISH data from this study and tested pairs from other studies. See Fig. S5 for plots including FlowFISH data only.

Using this data, we sought to identify generalizable rules to explain which enhancers regulate which genes in the genome. To do so, we compared various predictors to our experimental results by means of a precision-recall plot (Fig. 3A). (Precision refers to the proportion of positive predictions that are ‘true positives’ — where true regulatory connections are the 98 significant DE-G pairs where perturbation of the element led to a decrease in gene expression, and the 2912 non-regulatory connections are those where no decrease was detected despite >80% power to detect 25% effects. Recall refers to the proportion of true connections included in the predictions. For analysis of repressive effects, see Note S1).

**Fig. 3.**
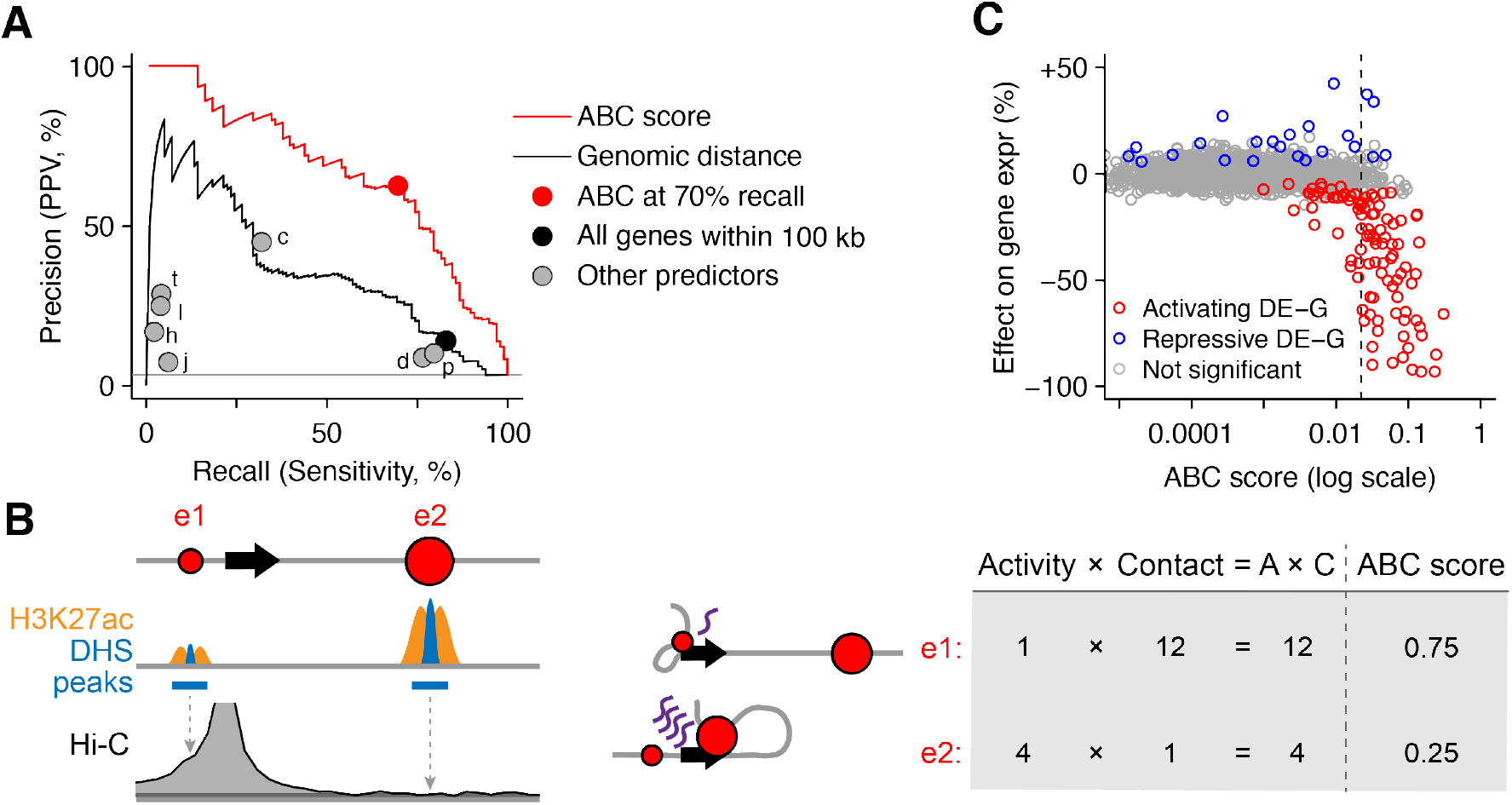
The ABC model predicts the target genes of enhancers. (**A**) Precision-recall plot for classifiers of DE-G pairs. Positive DE-G pairs are those where the distal element significantly decreases expression of the gene. Curves represent the performance for predicting significant decreases in expression for DE-G pairs based on thresholds on the ABC score (red) and genomic distance between the DE and the TSS of the gene (black). Circles represent the performance of various predictors in which DEs are assigned to: the closest expressed gene (“c”); all promoters within 100 kb (black), genes predicted by the algorithms TargetFinder (“t”) (*18*) or JEME (“j”) (*19*); promoters in same Hi-C contact domain (“d”); and promoters at the opposite anchors of Hi-C loops (“l”), RNA Polymerase II ChIA-PET loops (“p”) (*36*), or H3K27ac HiChIP loops (“h”) (*37*). (**B**) Calculation of the ABC score (see Methods). Values for DHS, H3K27ac, and Hi-C are presented in arbitrary units. (**C**) Comparison of ABC scores (predicted effect) with observed changes in gene expression upon perturbations. Each dot represents one tested DE-G pair. Dotted black line marks 70% recall, corresponding to the red dot in panel A.

We first examined three categories of methods that are commonly used to predict enhancer-gene connections, and found these had only modest predictive value (Fig. 3A):

1. Predictions based solely on distance thresholds along the genome performed poorly. For example, while 84% of regulatory DEs were located within 100 kb of their target promoter, only 14% of DEs within 100 kb of an expressed promoter had a regulatory effect (precision = 14%, recall = 84%). Assigning each DE to the closest expressed gene yielded 45% precision and 33% recall.
2. Predictions based solely on features of the 3D genome also performed poorly. Assigning each DE to promoters based on the presence of Hi-C loops yielded 25% precision and 4% recall, and assigning each DE to each other promoter in the same Hi-C contact domain yielded 14% precision and 77% recall.
3. Predictions based on prior machine learning approaches were similarly unsuccessful, including supervised methods to predict enhancer-promoter interactions from epigenomic data and unsupervised methods based on correlations between chromatin marks and gene expression across cell types (Fig 3A, see Methods) (*18, 19*).

Given the limitations of existing methods, we developed a new Activity-by-Contact (ABC) model to predict enhancer-gene connections. This model is based on the simple biochemical notion that an element’s quantitative effect on a gene should depend on its strength as an enhancer (“Activity”) weighted by how often it comes into 3D contact with the promoter of the gene (“Contact”), and that the *relative* contribution of an element on a gene’s expression (as assayed by the proportional decrease in expression upon CRISPR-inhibition) should depend on the element’s effect divided by the total effect of all elements. Under this model (Fig. 3B), the fraction of regulatory input to gene G contributed by element E is thus given by:

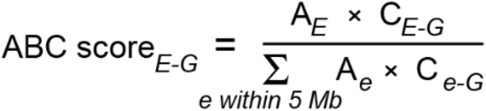

We defined Activity (A) as the geometric mean of the read counts of DHS and H3K27ac ChIP-Seq at an element E, and Contact (C) as the normalized Hi-C contact frequency between E and the promoter of gene G (see Methods). (The ABC score performed similarly across a range of data preprocessing parameters, and when defining Activity using other combinations of measurements of chromatin accessibility, histone modifications, and nascent transcription, see Methods, Fig. S6,S7,S8).

The ABC model performed remarkably well, and much better than alternatives, at predicting DE-G connections in our CRISPR dataset. The quantitative ABC score correlated with the experimentally measured relative effects of candidate elements on gene expression (Spearman *ρ* for regulatory DE-G pairs = –0.68 Fig. 3C). Binary classifiers based on thresholds on the ABC score substantially outperformed existing predictors of enhancer-gene regulation. For example, when we used an ABC threshold corresponding to 70% recall, the predictions had 63% precision, and the area under precision-recall curve (AUPRC) was 0.66, compared to 0.36 for predictions based solely on genomic distance (Fig. 3A).

The ABC score far outperformed models based on either Activity (quantitative DHS or H3K27ac signal) or Contact (Hi-C contact frequency) alone (AUPRC = 0.25, 0.17 and 0.31, respectively; Fig. 3A; Fig. S6A). This is because experimentally observed regulatory DE-G pairs varied substantially — with some having higher Activity and lower Contact, some showing higher Contact and lower Activity, and some having a balance of the two factors (Fig. S6B).

Given the ability of the ABC model to make predictions in K562 cells based solely on epigenomic data from that cell type, we explored whether the ABC model could generalize to predict enhancer-gene connections in other cell types.

To do so, we first identified alternative ways to estimate Contact in the ABC model; although maps of chromatin accessibility and histone modifications are available in many cell types, maps of 3D contacts are not. Because contact frequencies in Hi-C data correlate well across cell types (Note S3) (*38, 39*), we compared versions of the ABC model in which we estimated Contact for each DE-G pair using either K562 Hi-C data or the average Hi-C contact frequency from 8 other human cell types. Both approaches performed similarly at predicting our CRISPR data in K562 cells (AUPRC = 0.66 and 0.68 respectively; Fig. S9A). Thus, the ABC model can make predictions in a given cell type without cell-type specific Hi-C data, and minimally requires: (i) a measure of chromatin accessibility (DHS or ATAC-seq) and (ii) a measure of enhancer activity (ideally, H3K27ac ChIP-seq).

Using this approach, we evaluated the ability of the ABC model to predict 968 measured DE-G pairs in 5 additional human and mouse cell types beyond our initial K562 dataset (Table S3). These pairs included 940 from previous studies that inhibited DEs with epigenetic or genetic perturbations and measured the effects with RNA-seq or qPCR (*37, 40–49*), and 28 from new experiments in which we deleted enhancers in mouse embryonic stem cells and measured the effects using allele-specific RNA-seq (see Methods; Table S2). We used epigenomic datasets to generate genome-wide predictions of enhancer-gene connections in each of these 5 cell types (Table S6), and compared them to the CRISPR data in the corresponding cell type. The ABC scores correlated with the quantitative effects on gene expression (Spearman *ρ* for regulatory DE-G pairs = −0.38, Fig 4A), and at an ABC threshold corresponding to 70% recall, the predictions had 74% precision (AUPRC = 0.75, Fig 4B, see Methods). As expected, the predictions of the ABC model were highly cell-type specific: when we used ABC scores from K562 cells to predict DE-G pairs measured in other cell types, the AUPRC dropped from 0.75 to 0.12.

**Fig. 4.**
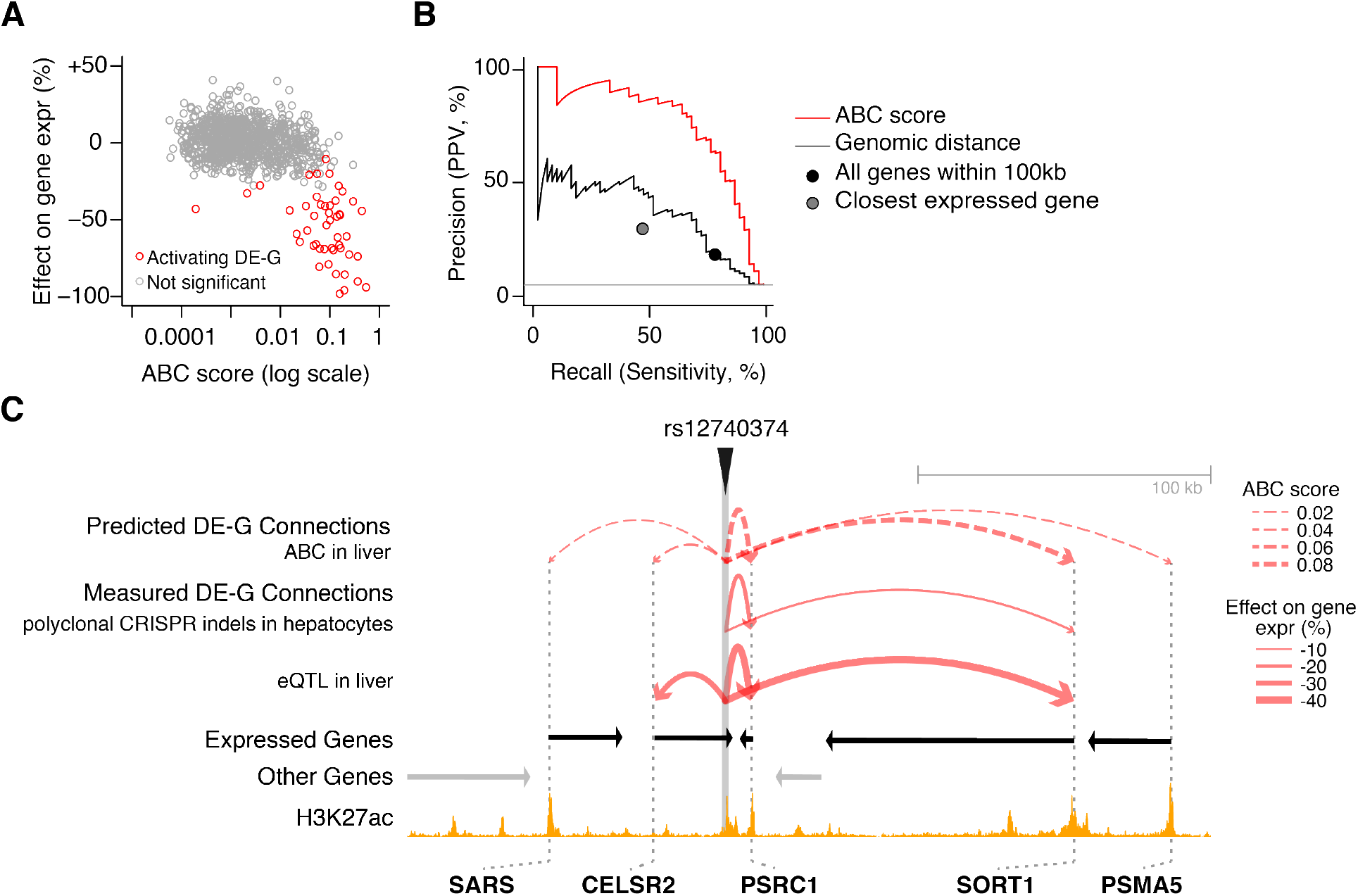
The ABC model generalizes across cell types. (**A**) Comparison of ABC scores (predicted effect) with observed changes in gene expression upon perturbations in GM12878 cells, LNCaP cells, NCCIT cells, primary human hepatocytes, and mouse ES cells. Each dot represents one tested DE-G pair. (**B**) Precision-recall plot for classifiers of DE-G pairs shown in (A). Positive DE-G pairs are those where the distal element significantly decreases expression of the gene. Curves represent the performance for predicting significant decreases in expression for DE-G pairs based on thresholds on the ABC score (red) and genomic distance between the DE and the TSS of the gene (black). Circles represent the performance of models that predict significant regulation for DE-G pairs based on various criteria: pair lies within 100 kb (black), and DEs are assigned to regulate the nearest expressed gene (purple). (**C**) Comparison of observed and predicted DE-G connections in the *SORT1* locus (chr1:109714926-109989926). Predicted DE-G connections (dotted red arcs) are based on ABC maps in primary human liver tissue. Observed DE-G connections (solid red arcs) are from previous experiments in which CRISPR was used to introduce indels near rs12740374 in primary hepatocytes (*49*) and an eQTL study in human liver (*50*).

We next examined the 16 DE-G pairs in our dataset that involved enhancers that harbor noncoding genetic variants known to influence risk for human diseases or traits. At a threshold corresponding to 70% recall in our K562 dataset, the ABC model correctly connected these DEs to their target gene(s) in 13 of 16 cases (81% recall). For example, a previous study identified 3 enhancers that contain noncoding variants associated with rare erythroid disorders, and found that introducing indels into these enhancers in K562 using CRISPR affected the expression of nearby genes involved in erythropoiesis (*33*). The ABC model correctly identified each of these 3 regulatory DE-G pairs in K562 cells, and, notably, also identified the same connections in primary human erythroid progenitor cells (Table S6). As another example, a variant associated with coronary artery disease and plasma low-density lipoprotein cholesterol (rs12740374) has been shown to be an eQTL for *SORT1* in primary human liver tissue, and CRISPR edits in the corresponding element affect *SORT1* expression in primary hepatocytes (*49, 50*). ABC maps in primary human liver tissue correctly connected this enhancer to *SORT1* (Fig. 4C). Thus, the ABC model can predict enhancer-gene connections based on cell-type specific epigenomic data, and may be widely useful for interpreting the functions of noncoding genetic variants associated with human diseases.

Finally, toward further improving predictions, we identified situations in which the ABC model failed to accurately predict DE-G connections.

We first compared predictions for tissue-specific versus ubiquitously expressed genes (sometimes referred to as “housekeeping” genes, see Methods), and found that the ABC model performed dramatically better for tissue-specific than for ubiquitously expressed genes (AUPRC = 0.77 vs 0.12). This was because ubiquitously expressed genes had fewer enhancers: for the 30 genes for which we had data for all nearby DEs, tissue-specific genes (n=22) had an average of 2.6 distal enhancers per gene, while ubiquitously expressed genes had only 0.1 (only a single enhancer across 8 ubiquitously expressed genes; rank-sum test *p* = 0.002, Fig. S10).

Interestingly, these observations are consistent with findings in *Drosophila*, where plasmid-based reporter assays have shown that ubiquitously expressed gene promoters are less sensitive to distal enhancers than are tissue-specific gene promoters (*6*). We conclude that the ABC model applies well to tissue-specific genes (97% of all genes, see Methods) but not to ubiquitously expressed genes, which appear to be largely insensitive to the effects of distal enhancer perturbations for reasons that remain to be explored.

We next examined our CRISPR dataset for DE-G pairs that likely represent regulatory effects due to mechanisms other than the cis-acting functions of enhancers (Note S1). We identified effects of distal CTCF sites, which may regulate gene expression by affecting 3D contacts (13 regulatory pairs, Fig. S11) and indirect effects, such as an enhancer regulating one gene that in turn affects a second nearby gene in *trans* (18 pairs, Fig. S12). Because these DE-G pairs do not represent direct effects of enhancers, we reasoned that removing them from the CRISPR dataset should provide a better estimate of the ability of the ABC model to predict enhancer-gene connections. Indeed, the AUPRC rose from 0.66 to 0.72 for all genes and to 0.82 for tissue-specific genes (Fig. S13). These results suggest a strategy to iteratively refine our predictions of DE-G connections by using CRISPRi tiling to identify exceptions to the ABC model, characterizing their molecular mechanisms, and developing new models to predict these effects. In summary, our work reveals key properties of enhancer-gene connections and provides an important foundation for future studies of regulatory elements and genetic variants in the noncoding genome. Our perturbation data, consistent with the predictions of the ABC model, indicate that enhancers often regulate more than one gene (Fig. 2D), that most enhancers with detectable effects are located within 100 kb of their target promoters (Fig. 2E), and that enhancers can have a wide range of quantitative effects on gene expression — including many elements with small effects (Fig. 3C).

Our results raise the intriguing possibility that the ABC model reflects an underlying biochemical mechanism: that enhancer specificity may often be controlled by quantitative factors including enhancer activity and enhancer-promoter contact frequency, rather than by qualitative logic involving the particular combinations of transcription factors at the enhancers and promoters. CRISPRi-FlowFISH and the ABC model provide a means to test these mechanisms, and to further refine our understanding of noncoding regulatory elements by mapping and modeling promoter-promoter regulation, functions of CTCF sites, and combinatorial effects of multiple enhancers in a locus.

Beyond its conceptual implications concerning gene regulation, the ABC model has important practical applications. Because it can make genome-wide predictions in a given cell type based on easily obtained epigenomic datasets, the ABC model provides a framework for mapping enhancer-gene connections across many cell types — including primary human cell types and states that are difficult to directly manipulate with CRISPR. This suggests a systematic approach to decode transcriptional regulatory networks and to interpret the functions of noncoding genetic variants that influence risk for human diseases and traits.

## Acknowledgements

We thank J. Chen, A. Chow, B. Cleary, C. de Boer, A. Dixit, M. Guttman, R. Herbst, C. Lareau, S. Rao, J. Ray, S. Reilly, R. Tewhey, J. Ulirsch, and C. Vockley for discussions and J. Marshall for assistance with fluorescent microscopy. FACS sorting was performed at the Broad Institute FACS Core by P. Rogers, S. Saldi, and C. Otis. This work was supported by funds from the Broad Institute (E.S.L.) and by NIH NHGRI Grant No. 1K99HG009917-01 to J.M.E. J.M.E. is supported by the Harvard Society of Fellows. S.R.G. is supported by National Institute of General Medical Sciences grant T32GM007753. E.S.L. serves on the Board of Directors for Codiak BioSciences and Neon Therapeutics, and serves on the Scientific Advisory Board of F-Prime Capital Partners and Third Rock Ventures; he is also affiliated with several non-profit organizations including serving on the Board of Directors of the Innocence Project, Count Me In, and Biden Cancer Initiative, and the Board of Trustees for the Parker Institute for Cancer Immunotherapy. He has served and continues to serve on various federal advisory committees. J.M.E., C.P.F., and E.S.L. are inventors on a patent application filed by the Broad Institute related to this work. Data presented in this paper can be found in the supplementary materials and in GEO accession GSE118912.

## Supplementary Notes

### Note S1. Additional mechanisms of distal regulatory elements

We considered two situations in which distal elements might have effects on gene expression through mechanisms distinct from or above that of enhancers: indirect effects and CTCF-bound elements. In addition to explaining some of the activating effects of distal elements (10 out of 98), these two situations also account for most of the DE-G pairs with repressive effects (20 out of 24).

#### Indirect regulatory effects of distal elements

The first situation involves indirect regulatory effects. For example, an enhancer that activates gene A might appear to repress B in the event that activation of A represses B. We noted that 24 of the 98 significant DE-G pairs (20%) in our data involve elements that, upon CRISPRi inhibition, led to *increased* expression (average +15%) of a nearby gene. However, these effects do not appear to result from cis-acting “repressors”; 6 of these 19 unique elements have activating effects on at least one other nearby gene (Fig. S12A). In one case, we verified that apparent repressive effects of an element on *PLP2* expression are due to that element activating *GATA1*, which in turn represses *PLP2* via a trans-acting function of the GATA1 protein product (Fig. S12B-D).

#### CTCF Sites

The second situtation involves elements bound by CTCF, a protein that affects gene regulation by shaping 3D genomic architecture (*51*) (34% of tested DHS elements bind CTCF). Notably, some CTCF sites appear to be coincident with enhancer elements (in that they are strongly marked by H3K27ac), while others appear to be separate. When we divided CTCF-bound distal DHS sites into H3K27ac^high^ vs. H3K27ac^low^ elements, we found clear differences between the two classes (Fig. S11). H3K27ac^high^ CTCF elements had larger effects on gene expression (average 32%) and were far more often activating rather than repressive (23 vs. 3), consistent with these elements primarily affecting gene expression as enhancers. The ABC model accurately predicts the effects of the perturbation of these elements (AUPRC = 0.53, Fig. S11B). In contrast, H3K27ac^low^ CTCF elements had smaller effects (average 10% vs. 32% for H3K27ac^high^ CTCF elements, rank-sum test *p* = 0.002), had balanced effects on gene expression (5 activating and 8 repressive vs. 23 and 3 for H3K27ac^high^ CTCF elements, Fisher’s exact *p* = 0.002), and the ABC model performed less well (AUPRC = 0.11, Fig. S11C).

### Note S2. Regulatory effects of promoters on nearby genes

In addition to DE-G pairs, our CRISPR dataset in K562 cells included 1114 distal promoter-gene (DP-G) pairs (where the CRISPR-targeted element is located <500 bp from a TSS).

We explored whether, beyond DE-G pairs, the ABC model did a good job of predicting DP-G connections – that is, regulatory effects of one promoter on the promoter of another nearby gene. In fact, it did not. Our dataset in K562 cells included 53 significant DP-G pairs (out of 1114 total tested), and the ABC score was only moderately predictive of these effects (AUPRC=0.16, Fig. S14). Importantly, the DP-G pairs in our dataset behaved qualitatively differently from the DE-G pairs: promoters more frequently had repressive effects (27 of 53 DP-G pairs, 51%, versus 20% for DE-G pairs, Fisher’s exact *p* < 10^−4^).

Promoters are known have the ability to affect the expression of neighboring genes through several mechanisms, including: activation of nearby genes in *cis*, for example by acting as an enhancer (*43, 52*); second-order, downstream effects of the promoter’s protein product; promoter-promoter competition, in which two promoters are proposed to compete for nearby regulatory elements (*53*); and transcriptional interference, in which transcription of one gene physically blocks transcription of another (*54*). We observe likely instances of each of these in our CRISPR dataset, detailed below.

#### *Cis* activation

We and others have shown that many gene promoters activate a neighboring gene in *cis* through DNA-mediated functions of their promoters (*43, 52, 55*). In this dataset, promoters that activated a nearby gene indeed had higher 3D contact with their target genes compared to other nearby genes (rank-sum *p* = 0.001).

#### Second-order *trans* effects

Effects on nearby genes observed when inhibiting a promoter may be second-order effects mediated by functions of the RNA or protein product, rather than first-order, *cis* effects of the promoter itself. We examined the 5 promoters whose inhibition affected 2 or more tested genes in our FlowFISH dataset (*GATA1, KLF1, LYL1, PPP1R15A*, and *SEC61A1*). Of these, 3 encode transcription factors and 2 encode regulators of translation, consistent with these genes having widespread effects on gene expression. For 3 of these genes, we found additional evidence to support that these effects on nearby genes did not result from direct *cis* effects of the promoter: inhibiting distal elements that regulate these genes had directionally consistent effects on other genes. These 5 promoters also more often had repressive effects than other promoters we found to affect the expression of nearby genes (median 2 repressed genes vs 0, rank-sum test *p* = 0.004). Based on this evidence, we expect that the effects of these 5 promoters on nearby genes are likely due to second-order, downstream effects of their protein products in *trans*.

For example, inhibiting the promoter of *GATA1* with CRISPRi led to increased expression of 3 nearby genes, and we confirmed through siRNA knockdown experiments that these effects are likely to result from *trans* functions of the GATA1 protein (Fig. S12D).

#### Promoter competition

In addition to acting through a *trans* function of its product, promoters may inhibit nearby genes by competing for enhancers or other activating signals. Our dataset included 18 promoters that appeared to repress a nearby gene. Notably, these included 2 promoters near *HBE1* and 1 near *MYC* that have been previously shown to compete with *HBE1* or *MYC* for activating signals in the genome (*56, 57*).

#### Transcriptional interference

We identified 4 promoters (2 alternative promoters for each of 2 genes) where CRISPR perturbation caused an increase (6-36%) in the expression of a convergently transcribed neighboring gene. In each of these cases, precision run-on sequencing (PRO-seq) showed that the transcriptional units of these genes overlap (Fig. S14C), suggesting that these promoters might repress the neighboring gene via transcriptional interference (*54*).

### Note S3. Alternative methods to estimate Contact in the ABC score

We explored alternative methods to estimate Contact in the ABC score in order to understand which features of genome architecture — such as loops and domains — are important for good prediction.

Because >70% of the variance in Hi-C contact frequencies across a chromosome can be explained by modeling chromatin as a featureless, uniform polymer in the condensed (globular) state (*58*) (see Methods), we tested simply using the theoretical contacts expected from extrusion globule and fractal globule models (Contact_*Globule*_ is proportional to Distance^−γ^, with γ = 0.7 and 1, respectively) (*58*). Both scores performed nearly as well as the ABC score based on Hi-C data (AUPRC = 0.64 for both, versus 0.68 for ABC, Fig. S9A,C). In comparison, Activity x Loop, Activity x Domain, Activity x Distance, and Activity x Contact_*Globule*_ models with more extreme values of γ performed less well (Fig. S9). These results show that the ABC model can predict DE-gene regulation reasonably well even without using information about locus-specific or cell-type specific features of the 3D genome. This yields a useful rule of thumb: 10-fold greater genomic distance between an enhancer and promoter leads to approximately 10-fold lower contact frequency and 10-fold smaller predicted effects on gene expression.

Notably, however, locus-specific Hi-C data did appear to yield better predictions for some DE-G pairs, including for long-range enhancer-gene connections in the *MYC* locus that coincide with the anchors of 3D loops (Fig. S9G,H). These and other 3D loops are present across many cell types (*17, 59*). Accordingly, we tested estimating Contact for a given pair of loci using the average contact frequency for those loci in Hi-C data from 8 other human cell types. We found that a Activity x Contact_*Average*_ model did a better job at predicting connections in the *MYC* locus than the Activity x Contact_*Globule*_ models, and had slightly better performance in the full K562 CRISPR dataset (AUPRC = 0.68 versus 0.64 respectively; Fig. S9A).

Together, these results indicate that cell-type specific features of the 3D genome are not required for good predictions, and that the relationship between genomic distance and quantitative contact frequency — more so than loops or domains — contains important information about regulatory enhancer-gene connections. These observations allow us to calculate ABC scores in a given cell type even without Hi-C data from that cell type.

## Methods

### Tissue Culture

We maintained K562 (ATCC) cells at a density between 100K and 1M per ml in RPMI-1640 (Thermo Fisher Scientific, Waltham, MA) with 10% heat-inactivated FBS (HIFBS, Thermo Fisher Scientific), 2mM L-glutamine, and 100 units/ml streptomycin and 100 mg/ml penicillin. We maintained HEK293Ts between 20 and 80% confluence in DMEM with 1 mM Sodium Pyruvate, 25mM Glucose (Thermo Fisher Scientific) and 10% HIFBS. CRISPRi-FlowFISH and qPCR experiments used K562 cells expressing KRAB-dCas9-IRES-BFP from a third generation tet-inducible promoter (Addgene # 85449).

### Individual gRNA qPCR

We generated stable cell lines expressing single gRNAs (Table S4) by lentiviral transduction in 8 μg/ml polybrene by centrifugation at 1200 x g for 45 minutes with 200,000 cells per well in 24 well plates. After 24 hours, we selected for transduction with 1 μg/ml puromycin (Gibco) for 72 hours then maintained cells in 0.3 μg/ml puromycin. For each gRNA, we generated 2 independent polyclonal cell populations through duplicate infections. We isolated RNA, made cDNA, and performed RT-qPCR as previously described (*17*) using primers listed in Table S4.

### Defining candidate elements

We defined candidate regulatory elements in 6 human cell types (K562, GM12878, NCCIT, LNCaP, primary hepatocytes, and primary erythroid progenitors), and 1 mouse cell type (mESCs).

For K562 and mESC, we concatenated all peaks called by ENCODE in both replicate DNase-seq experiments and merged resulting peaks (Table S5). This resulted in 174,403 peaks in K562. We then removed any peaks overlapping regions of the genome which have been observed to accumulate anomalous number of reads in epigenetic sequencing experiments (‘blacklisted regions’ (*60*) downloaded from https://sites.google.com/site/anshulkundaje/projects/blacklists) — with the exception of 5 peaks in mESCs, which were either tested by CRISPR experiments or which were promoters of genes nearby tested elements, and which were not removed. Given that the ENCODE peaks were initially 150bp in length, we extended each of these peaks 175bp to arrive at candidate elements that were 500bp in length.

For GM12878, NCCIT, LNCaP, primary hepatocytes, and primary erythroid progenitors we called peaks using MACS2 based on either DNase-seq or ATAC-seq as a measure of chromatin accessibility (Table S5). We initially considered all peaks with pvalue < .1 and removed peaks overlapping blacklisted regions. We then resized these peaks to be 500bp in length centered on the peak summit. In order to approximately match the number of candidate elements considered in K562, we then counted DNase-seq (or ATAC-seq) reads overlapping these regions and kept the 175,000 regions with the highest number of read counts. To this peak list, we added 1 kb regions centered on the transcription start site of all genes.

Any overlapping peaks resulting from this extension within a cell type were merged. We define these extended and merged peaks as *candidate elements*.

### Guide selection for CRISPRi-FlowFISH screens

We designed gRNAs within K562 candidate elements as previously described (*17*) and used all gRNAs within each candidate element after removing those with specificity scores <50 or with homopolymer stretches of more than 7 As, Gs, or Cs, or 4 Ts (Table S1).

### Gene selection for CRISPRi-FlowFISH screens

We used a series of filters for each probeset and screen to ensure robust, comprehensive, and quantitative discovery of regulatory elements for each gene (Fig. S4). We initially tested PrimeFlow probesets for genes expressed at >20 TPM in K562s in five genomic loci (Fig. S3). We first screened probesets by flow cytometry and selected those with >2-fold signal vs unstained cells. Next we performed a tiling CRISPRi-FlowFISH screen (see below) and focused our analysis on the screens that showed the following characteristics: (i) maximum unscaled knockdown among 20-gRNAs windows within 500 bp of the TSS >50%; (ii) variance in non-targeting, negative-control gRNAs <1; and (iii) >80% power to detect a 25% effect in at least 80% of elements (see below). Based on these filters, we performed and analyzed CRISPRi-FlowFISH screens for 28 genes.

### CRISPRi-FlowFISH Screens

We cloned gRNA libraries purchased from CustomArray (now GenScript) for each of 5 genomic loci (Fig S3), transduced into K562s harboring a doxycycline-inducible KRAB-dCas9, and selected for transduced cells as previously described (*17*). We induced KRAB-dCas9 expression with 1 μg/ml doxycycline for 48 hours. We used 30M cells for each screen.

We used the PrimeFlow RNA Assay Kit (Thermo Fisher; Catalog number: 88-18005) according to the manufacturer’s instructions with some modifications. Specifically, we split each screen into three 10 million cell reactions and performed five total washes with 35°C wash buffer after following the staining protocol. We stained each sample for the gene of interest with an Alexa Fluor 647 (AF647, “Type 1”) probeset and against a positive control housekeeping gene with Alexa Fluor 488 (AF488, “Type 4”). For most screens we used control gene *RPL13A*, but because BAX, *BCAT2, FTL, NUCB1*, and *PPP1R15A* are <700 kb from *RPL13A*, we used *ACTB* for these. Probeset used are listed in Table S4.

### Sorting

We diluted the stained cells in PBS with 0.5% BSA to a concentration of 2×10^7^ cells/ml and filtered using a 30μm filter (CellTrics, Catalog number 04-004-2326). We sorted 30 million cells for each screen into six bins based on fluorescence intensity of target genes using the Astrios EQ Sorter (Beckman Coulter B25982). To control for differences in staining efficiency for each cell, we normalized the fluorescence associated with the gene of interest to that of the control gene. Specifically, we used the color compensation tool to subtract a portion of each cell’s AF647 signal based on the intensity of its AF488 signal such that the mean AF488 signal in the top and bottom 25% of cells based on AF647 was within 10%. If necessary, we then reduced the level of compensation until the fraction of cells with AF647 signal equal to 0 was no more than 5%. We set the gates for each bin on the compensated signal to capture 10% of the cells according to the percentiles (i) 0-10% (ii) 10-20%, (iii) 35-45%, (iv) 55-65%, (v) 80-90%, and (vi) 90-100%.

### Genomic DNA extraction and gRNA sequencing

We collected the sorted cells by centrifugation at 800g for 5 minutes, resuspended cells in 100uL of Lysis buffer (50mM Tris-HCl, pH 8.1, 10mM EDTA, 1% SDS), and incubated at 65°C for 10 minutes for reverse crosslinking. Once the samples cooled to 37°C, we added 2ul of RNase Cocktail (Invitrogen, catalog #AM2286), mixed well, and incubated the mixture at 37°C for 30 minutes. Finally we added 10μl Proteinase K (NEB, catalog number P8107S), mixed well, and incubated the mixture at 37°C for 2 hours followed by incubation at 95°C for 20 min. We extracted genomic DNA using Agencourt XP (SPRI) beads (Beckman Coulter). We sequenced gRNA integrations as previously described (*17*).

### Analysis of CRISPRi-FlowFISH screens

To determine the effects of each gRNA on fluorescence, we used a maximum likelihood estimation (MLE) method. First, we normalized gRNA frequencies in each bin by dividing each gRNA count by the total read count for all gRNAs in that bin and summed normalized counts across PCR replicates. Next, we used the limited-memory Broyden–Fletcher–Goldfarb–Shanno (BFGS) algorithm MLE method in the R stats4 package to fit the read counts in each fluorescence bin to the log-normal distribution that would have most likely produced the observed counts in the bins. The effect size is from the mean of the log-normal fit. We assumed the gRNAs targeting the TSS of the assayed gene have a “true” effect size of 85% (based on previous observations that show CRISPRi effects of 80-90% across a panel of genes (*61*)), but that some portion of the FlowFISH signal is due to non-specific binding of the probe. Accordingly, we scaled the effect size of each gRNA within each screen linearly so that the strongest 20-gRNAs window within 500 bp of the target genes TSS gRNAs has effect size 85%. We then averaged the effect sizes of individual gRNAs across replicates.

To identify elements affecting the expression of the assayed gene, we used a *t*-test to determine whether the mean effect size of the gRNAs in each candidate element deviated significantly from the mean of scrambled-sequence, negative control gRNAs. We computed the FDR for elements using the Benjamani-Hochberg method applied per gene, and used an FDR threshold of 0.05 to call significant E-G interactions.

We excluded certain E-G pairs measured with CRISPRi-FlowFISH from further analysis (Table S2). E-G pairs were excluded if the pair met any of the below criteria:

i. There was less than 80% power to detect a 25% effect for this E-G pair.
ii. The element overlapped the gene’s promoter.
iii. The element was within the gene body or extended up to 2 kb downstream of the 3’ end of the gene.

### Enhancer perturbation data from other sources

To complement the data from our FlowFISH dataset, we curated results from previous experiments involving perturbations to accessible elements and precise measurements of the effects on gene expression. These included experiments involving a variety of perturbation methods (CRISPRi, 2-guide deletion, or other genome editing) and methods of measuring the effect on gene expression (RNA-seq, allele-specific RNA-seq, CRISPR screens, or RT-qPCR), and included six cell lines (K562, GM12878, NCCIT, LNCaP, hepatocytes, and mES cells). In cases where the same element-gene pair had been characterized in the same cell type by more than one group or by more than one assay, we included it only once in assessing the performance of the ABC model. We did not consider element-gene pairs where the element was that gene’s own promoter. Sources and study-specific details are annotated in Table S3. Additional details are included below, and in the following section (Power calculations).

#### Fulco 2016

We previously used CRISPRi (KRAB-dCas9) to tile gRNAs across a large region around *GATA1* and *MYC* in K562 cells and measured the effects using a proliferation assay (*17*). We used RT-qPCR data from this study to represent the effect sizes for the 7 and 2 enhancers that significantly affected *MYC* and *GATA1* expression, respectively. For all other elements, we estimated their effect sizes on gene expression based on the linear relationship between *MYC* expression and proliferation (*17*).

#### Klann 2017

Klann *et al*. used CRISPRi (dCas9-KRAB) to target gRNAs to DHS elements in a large region around *HBE1* in K562 cells and measured the effects by FACS sorting on an integrated HBE1-mCherry reporter (*23*). We downloaded the raw count file from this study (GSE96875) and filtered for gRNAs with a minimum total 50 reads across the high and low mCherry bins. We calculated the mean log2 fold-change across all replicates, and estimated effect sizes according to the linear relationship between this value and qPCR experiments for individual enhancers (Supp Figure 3B in Klann *et al*. 2017).

#### Ulirsch 2016

Ulirsch *et al*. used CRISPR used one gRNA per enhancer to introduce small deletions at each of 3 enhancers in K562 cells (*32*). We obtained the original qPCR data from the authors and assessed expression differences between homozygous knockout and wild-type clones using a *t* test.

#### Wakabayashi 2016

Wakabayashi *et al*. used one gRNA per enhancer to introduce small deletions at each of 5 enhancers in K562 cells (*33*). We obtained the original qPCR data from the authors and assessed expression differences between homozygous knockout and wild-type clones using a *t* test.

#### Thakore 2015

Thakore *et al*. used KRAB-dCas9 to inhibit an enhancer (HS2) in the globin locus in K562 and performed RNA-seq (*30*). We downloaded RNA-seq count matrices from GEO (GSE71557) and used DESeq2 to compute differential expression between biological replicate experiments using CR4 (the most effective guide RNA used in this study) versus no-guide controls. Genes within 1 Mb of the enhancer with FDR < 0.05 were considered true positives for downstream analysis; only genes within this range and with sufficiently high expression (>1 sample with read count >= 5) were considered in the multiple hypothesis correction.

#### Liu 2017

Liu *et al*. used KRAB-dCas9 to inhibit the promoters of several lncRNAs in K562 cells and performed RNA-seq (*34*). We downloaded the raw data from GSE85011 and quantified transcript abundance with kallisto (v. 0.43.0). A total of 19 RNA-seq experiments were performed; we removed one outlier (k562-LINC00910-1). We used DESeq2 to call differentially expressed genes for each of the 5 lncRNAs where two or more replicates were performed (EPB41L4A-AS1, LINC00263, LINC00909, MIR142, XLOC-042889). We compared the samples for a given promoter to all of the other samples (in which other lncRNA promoters were targeted) because there were no negative control samples. Genes within 1 Mb of the enhancer with FDR < 0.05 were considered true positives for downstream analysis; only genes within this range and with sufficiently high expression (>1 sample with read count >= 5) were considered in the multiple hypothesis correction.

#### Engreitz 2016

We previously generated homozygous and heterozygous knockout clones of 12 lncRNA and 6 mRNA promoters in mES cells on a 129S1/*Castaneus* hybrid genetic background, and measured the effects on gene expression using allele-specific RNA-seq (*43*) We calculated the average effects on the allelic expression of each gene within 1 Mb of the deleted promoter and included these in our perturbation database for this study. We assessed significance using DESeq2 to calculate the marginal effect of genotype (promoter knockout) after controlling for allele and sample (design formula = “~0 + Genotype + Allele + SampleName”). This effectively combines the allele-specific expression information across heterozygous and homozygous clones and leverages the statistical power of the empirical Bayes approach in DESeq2. We performed multiple hypothesis correction using the Benjamini-Hochberg method considering all genes within 1 Mb of the deleted promoter. This approach proved more powerful than the permutation-based method we previously used to analyze this data (*43*), and identified several additional nearby genes that showed significant allele-specific effects on expression. In Table S3 for this analysis, “nCtrl” and “nKO” refer to the number of wild-type and knockout *chromosomes* for each locus.

#### mES cell enhancer deletions (this study)

We also included data from new experiments in which we deleted two putative enhancers in mES cells via transfection of multiple gRNAs and measured the effects on nearby genes using allele-specific RNA sequencing, as previously described (*43*) (see Table S4 for gRNA and genotyping primer sequences). These two enhancers were selected on the basis of previous plasmid reporter assays showing enhancer activity for these elements (*62*) and are named “Chen2008-1” and “Chen2008-25” according to their number assignment from this previous study. We performed hybrid selection RNA-seq and produced allele-specific count tables as previously described (*43*). We assessed statistical significance using DESeq2 as described above. See Table S2.

#### Moorthy 2017

Moorthy *et al*. generated enhancer knockouts in mES cells on a 129S1/*Castaneus* hybrid genetic background, and measured the effects on gene expression using allele-specific RNA-seq as well as RT-qPCR (*46*). For the RNA-seq data, we calculated the average effects on the allelic expression of each gene within 1 Mb of the deleted element and assessed significance using DESeq2, considering allele-specific read counts in both heterozygous and homozygous clones as described above (*43*). This study generated a variety of heterozygous and homozygous deletions, including of multiple elements in different combinations in the same clones. We considered only the loci where at least one clone carried the deletion on the 129 allele and at least one clone carried the deletion on the *Castaneus* allele. For each deletion, we averaged the allele-specific effects across all clones. We looked for genes that showed >5% change in allele-specific expression with FDR < 0.25, but did not identify any significantly affected genes beyond those identified by the authors’ analysis.

#### Xie 2017

Xie *et al*. used KRAB-dCas9 and single-cell RNA-seq to identify 12 enhancers in K562 that significantly affect the expression of a neighboring gene (*29*). We used the log2 fold-change reported in the paper for genes whose expression was significantly affected by enhancer perturbations according to the authors’ analysis.

#### Blinka 2016, Huang 2018, Li 2014, Mumbach 2017, Rajagopal 2016, Spisak 2015, Tewhey 2016, Wang 2018, Xu 2015, Zhou 2014

For experiments from these studies, we estimated effect sizes and standard errors from figures in these studies, and assigned significance according to the authors’ analysis (*31, 35, 37, 40–42, 44, 45, 48, 49*).

#### Fuentes 2018

Fuentes et al. used CARGO to deliver an array of 12 gRNAs with dCas9-KRAB to simultaneously perturb LTR5HS, LTR5A, and LTR5B repeat elements (of which there are 910 annotated in the genome) in the NCCIT cell line, and measured the resulting changes in gene expression using RNA-seq (*47*). Because all elements were perturbed simultaneously (in each individual cell) in this study, the nature of the data is distinct from other data we analyzed, where only a single element was perturbed in any given experiment (or in any given cell in our CRISPRi screens). Accordingly, the data from Fuentes *et al*. required special analysis to identify DE-G pairs where effects on gene expression are likely to be due to the direct effects of an individual nearby DE/LTR.

We first identified the elements that were potentially targeted by Fuentes *et al*.: we considered 910 LTR5HS, LTR5A and LTR5B elements in the RepeatMasker (v4.0.5) database as well as 1194 dCas9 ChIP-Seq peaks (see below for ChIP-Seq analysis). We merged overlapping regions, resulting in 1427 candidate elements.

As different instances of the LTR5 repeats have high sequence similarity, we next determined how accurately we could measure the epigenetic profile (and thus the Activity compenent of the ABC score) of each LTR element. To determine the mappability of each element, we (i) simulated reads in each LTR region by tiling the region with 150bp paired-end reads of insert sizes between 150bp and 400bp (in increments of 10bp), (ii) mapped the simulated reads to the hg19 genome using BWA, and (iii) computed the fraction of reads from each LTR that map uniquely to that LTR (mapq >30). We considered the 1073 regions in which >95% of simulated reads mapped uniquely as sufficiently mappable for the purposes of the ABC score calculation.

In order to consider only the elements that were sucessfully perturbed in the CRISPRi condition, we further limited our analysis to the 1057 elements that displayed sufficient reduction in H3K27ac signal in the CRISPRi condition (>2-fold decrease in CRISPRi vs control condition, and less than 1 read per million in total H3K27ac ChIP-seq signal in the CRISPRi condition).

We next identified the set of genes that had exactly one nearby targeted LTR element (within 500 kb, not within the gene body). To assess changes in gene expression, we re-analyzed the RNA-seq data from Fuentes *et al*. (GSE111337): we quantified gene abundances using Kallisto (*63*) and computed differential expression with DESeq2 as described in Fuentes *et al*. (*47*). We considered a gene significantly differentially expressed if its Benjamini-Hochberg adjusted pvalue was <0.05. We calculated the statistical power to detect effects as described in the following section.

In order to reduce the contribution of *trans* effects, we applied a filter similar to that described in Fuentes *et al*. (*47*): we limited our analysis to genes that have concordant effects in the CRISPRi and CRISPRa conditions. Specifically, we only analyzed genes that were significantly down-regulated in the CRISPRi condition and up-regulated in the CRISPRa condition, or genes that were not significant in both conditions and that had sufficient power in both conditions.

To summarize, we applied the following to filters to the dataset generated by Fuentes *et al*:

We only considered LTR elements which

- Had sufficient decrease in H3K27ac signal upon CRISPRi perturbation
- Had sufficiently high simulated mappability
- Were at least 500kb from the closest other LTR element.
- Did not overlap a gene promoter

We only considered genes which

- Did not have an LTR within the gene body.
- Had concordant effects under perturbations by CRISPRi and CRISPRa
- Had exactly one LTR within 500kb

This resulted in a set of 22 positive and 872 negative LTR-gene pairs at the lenient power threshold (see below), and 22 positive and 0 negative LTR-gene pairs at the stringent power threshold (Table S3). We additionally considered 5 LTR-gene pairs where Fuentes *et al*. deleted the LTR and quantified the effect on the target gene by qPCR. The deletion of the LTR proximal to *EPHA7* was not included as this LTR element did not have sufficiently high simulated mappability.

### Power calculations for differential expression

Enhancers are known to have a wide range of effect sizes on gene expression (including examples as low as 10%) (*17*), and so we designed our experimental and computational analysis of enhancer-gene connections to precisely estimate effect sizes and carefully estimate the power to detect certain effect sizes. For all datasets (including in our FlowFISH data and from other sources), we assigned each tested element-gene pair into one of four categories: (i) statistically significant decrease on gene expression (“positive” for precision-recall analysis); (ii) statistically significant increase on gene expression (“negative” for precision-recall analysis); (iii) >80% power to detect a 25% effect on gene expression, but no significant effect detected (“negative” for precision-recall analysis); or (iv) <80% power to detect a >25% effect on gene expression (not considered in our analysis of element-gene connections due to lack of power). As this stringent power cutoff permited only 23 negative DE-G pairs for analysis of the perturbation data in other cell types (Fig. S15), we also tested using a lenient threshold of >80% power to detect a 50% effect on gene expression (Fig. 4), which increased the number of negative pairs in other cell types to 920.

#### Power calculations for FlowFISH experiments

For each candidate element, we used a t-test (equal variances) to compare the MLE effects of the gRNAs in that element to the MLE effects of 668-3505 negative controls (non-targeting gRNAs), and applied the Benjamini-Hochberg correction across the set of tests in each screen. We used summary statistics from these experiments (standard error of the mean and *n* for cases and controls) to analytically solve for the power to detect >25% changes in gene expression. We removed screens without 80% power to detect a 25% effect in at least 80% of elements, and additionally a single tested E-G connection with insufficient power.

#### Power calculations for qPCR datasets

We used a t-test (equal variances) to evaluate differences in gene expression for RT-qPCR datasets. We used summary statistics from these experiments (standard error of the mean and *n* for cases and controls) to analytically solve for the power to detect >25% or >50% changes in gene expression. *P*-value cutoffs for power calculations were determined using the multiple hypothesis correction methods used in the original studies.

#### Power calculations for RNA-seq datasets

We used DESeq2 to calculate differences in gene expression between cases (enhancer perturbation) and controls (*64*). DESeq2 uses a series of empirical Bayes steps to estimate the mean, variance, and log-fold-change for each gene. We cannot compute the power for this test analytically and instead used a simulation-based procedure to estimate the power to detect changes in the expression of each gene in each enhancer perturbation:

1. We considered the real RNA-seq data for each test, for example consisting of several replicates of case and control conditions.
2. We removed genes where fewer than two samples had five or more reads.
3. We estimated the mean and dispersion parameters for each gene using the DESeq2 empirical Bayes procedure.
4. Based on these parameters, we simulated 100 random datasets across all genes with the same total read counts as the original experiments. For each gene within 1 Mb of the perturbation, we reduced the mean parameter by 25% or 50% for these simulations.
5. We used the DESeq2 pipeline on each simulated dataset to compute the *p*-value for every gene in the genome. For each gene within 1 Mb of the perturbation, we computed the FDR by performing multiple hypothesis correction with the Benjamini-Hochberg method using the *p*-value of each gene in the simulated dataset together with the *p*-values of other genes within 1 Mb derived from the real data.
6. We computed power based on the fraction of the 100 simulations in which FDR < 0.05.

We used an identical procedure for power calculations for allele-specific RNA-seq, with the only difference being the inclusion of additional variables (representing allele and sample) in the DESeq2 design matrix.

#### Computing the effects of large deletions

In some cases, certain genomic perturbations (e.g., from Moorthy *et al*. 2017) involved large genomic deletions that spanned multiple ABC model elements. In these cases, we predicted the effect of the deletion as the sum of the ABC score of all overlapping elements, and assigned it to the “distal promoter” category if it overlapped a promoter element.

#### Stringent and lenient power filters for data in other cell types

We analyzed the enhancer perturbation data collated in other cell types at two different power thresholds, the “stringent” threshold we used for analysis of the K562 data (80% power to detect 25% effects on gene expression), and a “lenient” threshold of 80% power to detect 50% effects on gene expression because the experiments in other cell types were not as well powered as our CRISPRi-FlowFISH method, and thus assigned fewer non-regulatory DE-G pairs.

In the stringently-filtered dataset, applying the threshold on the ABC score corresponding to 70% recall and 63% precision in our initial K562 dataset could identify DE-G connections in other cell types with 91% recall and 75% precision (Fig S15).

When we relaxed the power requirements for data in other cell types to include more non-regulatory DE-G pairs (from 80% power for detecting 25% effects to detecting 50% effects), we found that the ABC model performed similarly in the K562 and cross-cell-type datasets (AUPRC = 0.66 vs 0.75, respectively; Fig 4).

### Epigenomic datasets, processing, and analysis

#### DNaseI hypersensitivity sequencing (DHS), ChIP-seq, and Expression datasets

We downloaded bam files for DNase I hypersensitivity sequencing (DHS), ChIP-seq for several chromatin marks including H3K27ac, and several transcription factors from a varienty of sources including ENCODE (see Table S5) (*47, 65–68*). We generated our own H3K27ac ChIP-seq data in F1 129/Castaneus hybrid mESCs grown in 2i media as previously described (*43*), and our own ATAC data in NCCIT cells as described below (available from GSE118912).

#### Hi-C

We analyzed K562 and GM12878 *in situ* Hi-C maps described previously (GSE63525) (*59*). We also generated new *in situ* Hi-C maps of male mouse V6.5 embryonic stem cells grown in 2i conditions as previously described (*59*), and sequenced 4 technical replicates to a combined depth of 1.17 billion reads (available from GSE118912). Hi-C loop and contact domain annotations were computed using the Juicer suite of tools (*69*).

#### NCCIT ATAC

We performed ATAC-seq on 10K cells NCCIT cells in duplicate according to the protocol described by Buenrostro *et al*. (*70*) with some modifications. Specifically, we used Sigma Nuclei EZ lysis buffer for lysis for 10 minutes while spinning 500xG at 4C, resuspended with the lysis buffer, and spun again for 3 minutes. We then resuspended the nuclei pellet with a tagmentation buffer containing 12.5 uL of TD buffer, 1.25 uL of Tn5 transposase, 7.5 uL of PBS and 2.75 uL of water. After 15 cycles of PCR we cleaned the products with Agencourt XP (SPRI) beads and sequenced to a depth of at least 30M reads per sample with 100 and 200 bp paired-end reads on a HiSeq 2500.

#### NCCIT ChIP-seq processing

For analysis of CRISPRi and CRISPRa data from Fuentes *et al*., we downloaded dCas9-GFP ChIP-seq data from GSE111337 and obtained H3K27ac ChIP-seq data directly from the authors (*47*). We aligned reads using BWA (v0.7.17) (*71*), removed PCR duplicates using the MarkDuplicates function from Picard (v1.731), and removed reads with mapq < 30. We used MACS2 (v2.1.1) (*72*) to call peaks on Cas9 ChIP-seq using the non-targeting conditions as controls as described in (*47*).

### Activity by Contact (ABC) model

We designed the Activity by Contact (ABC) score to represent a mechanistic model in which enhancers contact target promoters to activate gene expression. In a simple conception of such a model, the quantitative effect of an enhancer depends on the frequency at which it contacts a promoter multiplied by the strength of the enhancer (i.e., the ability of the enhancer to activate transcription upon contacting a promoter) (*17*). Moreover, the relative contribution of an element on a gene’s expression (as assayed by the proportional decrease in expression upon CRISPR-inhibition) should depend on the element’s effect divided by the total effect of all elements.

To extend this conceptual framework to enable computing the quantitative effects of enhancers on the expression of any gene, we formulated the ABC score:

> ABC score for effect of element E on gene G = Activity of E × Contact frequency between E and G / Sum of (Activity × Contact Frequency) over all candidate elements within 5 Mb.

Operationally, Activity (A) is defined as the geometric mean of the read counts of DHS and H3K27ac ChIP-seq at an element E, and Contact (C) as the normalized Hi-C contact frequency between E and the promoter of gene G, and elements are defined as ~500bp regions centered on DHS peaks.

This model has the following characteristics or assumptions:

1. The effect of an element on gene expression is linearly proportional to contact frequency and enhancer Activity.
2. A given enhancer has equal “Activity” for all genes — that is, it does not model the potential for biochemical specificity that could allow certain enhancers to regulate only certain promoters.
3. Different enhancers contribute additively and independently to the expression of a gene.
4. The sum in the denominator includes the gene’s own promoter, which is considered a potential enhancer with Activity calculated in the same manner as other enhancers.
5. The model computes the relative effect of an enhancer on gene expression, but does not estimate the absolute effect.
6. The model aims to predict the functions of enhancers, but not the functions of elements that act through other mechanisms.

We detail the calculation of the ABC score and discuss these assumptions below.

#### Calculating enhancer activity from DHS and H3K27ac ChIP-seq signals

We estimated enhancer activity of candidate elements using a combination of quantitative DNase-seq and H3K27ac ChIP-seq signals. For a given element, we counted DHS and H3K27ac reads (per million) in DNase peaks (150 bp from ENCODE), which we extended by 175 bp on either side (to 500 bp total; average length after merging overlapping peaks = 597 bp, Fig. S2B) because H3K27ac ChIP-seq signals are strongest on the nucleosomes flanking the nucleosome-free DHS peak. We computed the geometric mean of DNase-seq and H3K27ac ChIP-seq signals because we expect that strong enhancers should have strong signals for both, and that elements that have only one or the other likely represent other types of elements. (Elements with strong DNase-seq signal but no H3K27ac ChIP-seq signal might be CTCF-bound topological elements. Elements with strong H3K27ac signal but no DNase-seq signal might be sequences that are close by to strong enhancers but do not themselves have enhancer activity, due to the spreading H3K27ac signal over hundreds to thousands of bp.)

We note that this calculation of enhancer activity is the same for a given element across all genes. This means that the model assumes that an enhancer has the same “Activity” for every promoter (*i.e*., no differences due to biochemical specificity).

#### Calculating contact frequency from cell-type specific Hi-C data

In our initial analysis in K562 cells, we obtained the Contact component of the ABC score for E-G pairs from Hi-C data in K562 cells, using the quantitative signal observed in the 5-kb x 5-kb bin containing the center of E and TSS of G.

Specifically, we used KR-normalized Hi-C contact maps at 5-kb resolution, and processed these maps in two steps:

i. Rows and columns corresponding to KR normalization factors less than .1 were removed (these typically correspond to 5-kb bins with very few reads).
ii. Each diagonal entry of the Hi-C matrix was replaced by the maximum of its four neighboring entries. Justification: The diagonal of the Hi-C contact map corresponds to the measured contact frequency between a 5 kb region of the genome and itself. The signal in bins on the diagonal can include restriction fragments that self-ligate to form a circle, or adjacent fragments that re-ligate, which are not representative of contact frequency. Empirically, we observed that the Hi-C signal in the diagonal bin was not well correlated with either of its neighboring bins and was influenced by the number of restriction sites contained in the bin.

We then computed Contact for an E-G pair by rescaling the data as follows:

i. We extract the row of the processed Hi-C matrix that contains the TSS of G. For convenience, the row is rescaled so that the maximum value is 100.
ii. We set the Contact of the E-G pair to the Hi-C signal at the bin of this row corresponding to the midpoint of E.
iii. We add a small adjustment (“pseudocount”) to ensure that the contact frequency for each E-G pair is non-zero. For E-G pairs within 1 Mb, the adjustment is equal to the expected contact frequency at 1Mb (as predicted by the power-law relationship between contact frequency and genomic distance, see below), and for E-G pairs at distance d (d > 1Mb), the adjustment is equal to the expected contact at distance d. In each case the adjustment is scaled to be in the same units as described in (i). Adding the adjustment sometimes results in a quantitative Contact greater than 100; in such cases, the Contact is reduced to 100.

#### Calculating the contribution of one candidate element relative to others in the region

To calculate the relative effect of each element to the expression of a gene, we normalize the Activity by Contact of one element for a given gene to the sum of the Activity by Contact of other nearby elements. We included all elements within 5 Mb of the gene’s promoter in this calculation, and found that the performance of the model was not sensitive to this parameter (see below). We also included each gene’s own promoter as an element in the denominator of the ABC score. This is because the promoters of genes are known to have the potential to act as enhancers for other genes and are frequently bound by activating TFs (*43, 52*). Thus, the ABC score considers that the element near the TSS can have enhancer activity that contributes to the total regulatory signals relevant for that gene. We note that this normalization encodes the simplifying assumption that each element contributes independently and additively to gene expression. Based on the performance of the model in distinguishing significant DE-G pairs, this assumption appears sufficient for practical performance of the model. This first-order ABC model provides a foundation for incorporating higher-order effects such as the potential for nonlinear effects of multiple enhancers in a locus.

#### Sensitivity of the ABC score to chosen parameters

An attractive feature of the ABC model is its simplicity: at its core, the formula involves counting reads in DHS, H3K27ac, and Hi-C experiments, and performing a few addition and multiplication operations. We designed this ABC model based on the conceptual model of enhancer function described. Notably, there are no free parameters that need to be fit. While the model contains no free parameters, there are certain choices that need to be made in data processing. We made these choices based on known properties of epigenomic datasets. Specifically:

- We set the size extension of DHS peaks to 175 bp to include the nucleosome signal neighboring the DHS peak, and, together with the 150-bp size DHS peaks in ENCODE data, to yield extended elements with a convenient size (500 bp).
- We chose a genomic distance cutoff of 5 Mb based on this including all confirmed cases of *cis* regulation by enhancers — the longest of which is ~2 Mb.
- We regularized the Hi-C data by adding an adjustment factor (“pseudocount”), equal to the average contact at d = 1 Mb (as described above).
- We included the promoter of each gene as a regulatory element and assigned its “Contact” (with itself) according to the diagonal Hi-C signal as described above.

To determine if the performance of the ABC score was sensitive to these choices, we varied the size of extension of DHS peaks (range: 0 to 1000 bp; our choice was 175 bp), the genomic distance over which elements were included in the model (range: 500 kb to 10 Mb; our choice was 5 Mb), the Hi-C adjustment factor (range: average signal at 100 kb to 10 Mb; our choice was 1 Mb), and the signal at the diagonal bin of the Hi-C matrix relative to its neighboring bins (range: 0 to 500%; our choice was 100%). A broad range of parameter choices gave nearly identical performance (Fig. S8). The parameter that appeared most important was the size extension of DHS peaks, where either much lower or much higher extensions led to somewhat lower accuracy. This appears to be because at lower extension values, the H3K27ac signal is not properly captured, while at higher values the merging of nearby elements results in poor ability to distinguish between the functions of adjacent DHS peaks. These observations suggest that the ABC score is robust to our initial choices in data processing.

### Alternative methods to estimate Contact in the ABC model

#### Approximating Hi-C contact frequency with the average Hi-C data

To evaluate the performance of the ABC model using a non-cell-type-specific Hi-C dataset, we generated locus specific Hi-C profiles from an average of 8 human Hi-C datasets (Table S5). These averaged profiles were created as follows:

i. For each gene in the genome, we extract the row corresponding to this gene from each Hi-C matrix (KR normalized, at 5KB resolution)
ii. Each of these profiles is then scaled using the cell-type specific power law parameters relative to the K562 power law parameters (see below)
iii. Finally, the total Hi-C signal in each cell-type specific profile is normalized to sum to one and then averaged across cell types to create the average profile at a given locus

#### Normalizing Hi-C Profiles Using the Power-Law Fit

We find that different Hi-C datasets have slightly different power-law parameters. To weight all cell types equally in generating an average Hi-C profile, we scale the Hi-C profile in a given cell type by the cell-type specific parameters from the power law relationship in that cell type (see below). The scaling factor at distance *d* is given by (*scale*_ref_/*scale*_celltype_) * *d* ^ (*gamma*_ref_ − *gamma*_celltype_), where *scale*_ref_ and *gamma*_ref_ are the given reference parameters. For this study we used the power-law parameters in K562 as a reference.

#### Fitting a power-law relationship to Hi-C data

We fit a power-law relationship to the Hi-C data in a given cell type as follows:

i. We aggregate all entries of the Hi-C matrix located greater than 10kb and less than 1Mb from all gene promoters (KR normalized at 5kb resolution)
ii. We then perform a linear regression of the Hi-C signal in these bins on genomic distance in log-log space. The slope of this line is the *gamma* parameter and the intercept is the *scale* parameter.

#### Approximating Hi-C contact frequency with polymer globule models

To compute the variance in Hi-C contact frequencies (KR-normalized contacts) explained by a polymer globule model (and relevant to enhancer-gene regulation), we examined all gene TSSs and their contacts with loci at distances between 10 kb and 5 Mb in K562. The fractal globule model explained 69% of the variance in Hi-C contact frequency and the extrusion globule model explained 71% of the variance.

### Comparison of ABC predictions across cell types

#### Quantile normalization of epigenomic data

In order to facilitate a comparison of epigenomic datasets across cell types (and across assays, e.g., DNase-seq vs ATAC-seq), we quantile normalized the read counts in candidate elements from other cell types to the read counts in the corresponding assays in K562. Specifically, for each data type (H3K27ac ChIP-seq and DNase-seq or ATAC-seq) and for each class of element (promoter-proximal and distal), we quantile normalized the signal (in RPM) from this data-type and enhancer-class to the signal in K562. We then computed genome-wide ABC scores using these normalized epigenomic profiles as described above. If Hi-C data was not available in the cell type, we used the average Hi-C profile described above.

#### Identifying expressed genes for ABC predictions

When using the ABC model to predict enhancer-gene connections genome-wide (Table S6), we made predictions only for genes that are “expressed”. For cell types where RNA-seq data was available, we defined expressed genes as those with RNA-seq transcripts per million (TPM) > 1. For cell types where RNA-seq data was not available (LNCaP, primary liver) we defined expressed genes as those whose promoters had chromatin states consistent with active transcription. Specifically, we calculated a promoter score as the sum of DHS (or ATAC-seq) reads and H3K27ac ChIP-seq reads on a 1 kb region centered at the gene’s transcription start site, and then defined expressed genes and those with the top 60% of promoter scores.

### Comparison to other published enhancer-gene prediction methods

We evaluated the performance of the following published enhancer-gene prediction methods in predicting DE-G connections in our dataset:

JEME enhancer-gene predictions from from Cao *et al*. 2017. The Joint Effects of Multiple Enhancers (JEME) method first computes correlations between gene expression and various enhancer features (e.g., DNAase1, H3K4me1) across multiple cell types to identify a set of putative enhancers. Then a sample-specific model is used to predict the enhancer gene connections in a given cell type (*19*). We downloaded the lasso-based JEME predictions in K562 (ID 121) from http://yiplab.cse.cuhk.edu.hk/jeme/ and used the reported score as a continuous predictor.

K562 ChIA-PET loops from Li *et al*. 2012. We downloaded the K562 saturated PET clusters from Supplementary Table 2 of https://www.ncbi.nlm.nih.gov/pmc/articles/PMC3339270/#SD1 (*36*). For each E-G pair in our dataset, we searched to see if the element and gene TSS overlapped two interacting regions listed in this file. If so, the pair received a score of 1, otherwise it received 0.

TargetFinder enhancer-promoter predictions from Whalen *et al*. 2016. We downloaded the TargetFinder K562 predictions from https://github.com/shwhalen/targetfinder (*18*). We used the GBM classifier including Enhancer and Promoter windows (EPW). For each DE-G pair in our dataset, we searched to see if the element and gene TSS overlapped an enhancer and promoter loop listed in this file. If so, we assigned the pair a score corresponding to the ‘prediction’ column from this file, otherwise it received 0.

Hi-ChIP loops from Mumbach *et al*. 2017. We downloaded the HiCCUPS high-confidence loop calls from K562 cells from supplementary table 2 of https://www.nature.com/articles/ng.3963#supplementary-information (*37*). For each DE-G pair in our dataset, we searched to see if the element and gene TSS overlapped a loop listed in these files. If so, we assigned the pair a score of 1, otherwise it received 0.

### FlowFISH to study enhancers and promoters in the *MYC* locus

In our previous study, we identified 7 *MYC* enhancers that quantitatively tuned *MYC* expression (by 9-60%) (*17*). We studied the effects of these 7 enhancers on two other genes in the locus (*PVT1* and *CCDC26*, both noncoding RNAs) to examine the potential for these enhancers to specifically regulate certain genes. To test their effects in the genome, we designed a pool containing 2-3 gRNAs per gene and 13 negative control gRNAs. We used CRISPRi-FlowFISH for *MYC, PVT1*, and *CCDC26* to measure the effects of these 7 enhancers on the expression of each of these genes (Fig S9H. Because *PVT1* has multiple promoters in K562 cells (*73*), we verified the effects we observed in FlowFISH using RT-qPCR with primers corresponding to a specific *PVT1* isoform (that uses e3 as a promoter) as previously described (*17*).

### siRNA-mediated knockdown of GATA1

We transfected 200,000 K562 CRISPRi cells (from the same population of cells that was used in the CRISPRi-FlowFISH screens) with siRNAs (from Ambion, Thermo Fisher Scientific, Table S4) using the Amaxa Nucleofector 96-well Shuttle (Lonza, program: 96-FF-120) following the manufacturer’s protocol. We transfected each siRNA in quadruplicate. We harvested cells in buffer RLT (Qiagen, Germantown, MD) 48 hours after transfection and estimated target gene expression relative to cells transfected with non-targeting siRNAs by RNA sequencing.

For RNA-seq, we followed version 2 of a 3’ cDNA-enriched bulk RNA barcoding and sequencing (BRB-seq) protocol (*74*) with some modifications. We isolated RNA from 100,000 cells in RLT with 2.2X volume Agencourt RNAClean XP SPRI beads (Beckman Coulter, Danvers, MA). As measured by the RNA Qubit High Sensitivity Kit, we used 125 ng RNA input per sample during first strand synthesis with a barcoded RT primer. We then pooled 7-12 barcoded first-strand cDNA samples together. After an overnight second-strand synthesis using nick translation as previously described (*74*), we split each pool (containing multiple samples indexed during first strand synthesis) into 4-8 tagmentation replicates. We tagmented 5 ng of cDNA using 1 uL Nextera Tagment DNA Tn5 transposase (Illumina, San Diego, CA, 15027916) in a 10 uL tagmentation mix for 10 minutes at 55 °C.

Using the custom P5 primer and a standard Nextera i7 indexing primer, we used qPCR to optimize the number of PCR amplification cycles by chosing the cycle number that produced half the maximal fluorescent signal. We cleaned up the reaction twice using 0.8X volume Agencourt Ampure XP SPRI solution (Beckman Coulter, Danvers, MA). We sequenced the resulting libraries on a HiSeq 2500 (Illumina) with 35 bp reads.

We trimmed reads using BRB-seqTools v1.3, aligned reads to hg19 using STAR (v2.5.2b), and used BRB-seqTools v1.3 to count UMIs in RefSeq gene exons. We used DESeq2 to compute differential expression of siRNAs against *GATA1* versus non-targetting controls with the design formula “~perturbation+dose” (to control for the doses of siRNAs). Genes within 1 Mb of *GATA1* with Benjamani-Hochberg-corrected p-value < 0.05 were considered differentially regulated; only genes within this range and with sufficiently high expression (>1 sample with read count >= 5) were considered in the multiple hypothesis correction.

### Analysis of ubiquitously-expressed genes

To define the set of ubiquitously-expressed genes for human, we intersected 4 published lists of ubiquitously expressed genes from studies enumerating genes with detectable (*75*) or uniform expression across many tissues (*73, 76*) for 847 total ubiquitously expressed genes (Table S7). For mouse, we used the list of 4781 uniformly expressed genes provided in Li *et al*. (*77*). We refer to all other genes as “tissue-specific”.

To compute the number of enhancers per tissue-specific or ubiquitously-expressed gene, we focused on the subset of our data where we had comprehensive CRISPRi tiling data testing all elements near a genes, including 28 genes from this study and 2 genes (*MYC* and *HBE1*) from previous studies (*17, 23*). In this subset of the data, we found 58 regulatory DE-G pairs for the 22 tissue-specific genes and 1 regulatory DE-G pairs for the 8 ubiquitously-expressed genes (Fisher’s exact *p* < 10^−4^), as reported in the main text.

We note that the same trends hold in the full CRISPR dataset across all cell types (including DE-G pairs where we do not necessarily have comprehensive mapping of all DEs for that gene): we find more significant regulatory DE-G pairs for tissue-specific genes (140 significant pairs out of 2304 tested) than for ubiquitously expressed genes (6 significant pairs out of 777 tested, Fisher’s exact *p* < 10^−11^).

### Analysis of CTCF sites

We considered that CRISPRi perturbation of CTCF-bound elements may affect gene expression through effects on 3D genome contacts rather than that through disruption of enhancer elements. We downloaded CTCF ChIP-seq peak calls generated by ENCODE (Table S5) and labeled a distal element as a CTCF-bound if the element overlapped a CTCF ChIP-seq peak. We further classified each CTCF site as H3K27ac^High^ or H3K27ac^Low^, corresponding to elements with H3K27ac signal above or below the median H3K27ac signal for all tested distal elements in K562s.

### Estimating the performance of the ABC score at predicting enhancer-gene connections

To estimate the performance of the ABC score on a dataset measuring only the direct *cis*-effects of enhancers, we removed 762 total DE-G pairs that involved (i) CTCF-bound elements unlikely to function as enhancers (H3K27ac^Low^, 755 DE-G pairs), or (ii) DE-G pairs likely to result from indirect effects (18 DE-G pairs).

The latter category was defined as follows: We first identified genes (A) where the effects of promoter inhibition on nearby genes (G) are likely to be explained by second-order, indirect effects of the protein product (as described above). Enhancers that regulate gene A may also have indirect effects on gene G. Accordingly, we removed the 18 DE-G pairs where the element activates gene A and also affects gene G in a direction consistent with effect of promoter A on gene G.

The performance of the ABC score is markedly higher on this filtered dataset, with the AUPRC rising from 0.77 to 0.82 for tissue-specific genes and 0.66 to 0.72 for all genes (Fig. S13B). We note that all analyses presented in the paper use the full, unfiltered dataset in K562 cells unless otherwise specified.

### Software for data analysis and graphical plots

We used the following software for data analysis and graphical plots: R R (3.1.1) with Bioconductor (3.0)(*78*), Python (3.4.2), matplotlib (1.5.3), numpy (1.15.2), Pandas (0.23.4), Pybedtools (0.7.8), pyBigWig (0.3.2), pysam (0.13), scikit-learn (0.18.2), scipy (0.18.1), seaborn (0.7.1).

### Genome build

All coordinates in the human genome are reported using build hg19, and all coordinates in the mouse genome are reported using build mm9.

## Supplementary Tables and Figures

**Table S1 (separate file)**

**CRISPRi-FlowFISH library gRNA sequences**

Sequences and genomic coordinates of gRNAs used in CRISPRi-FlowFISH screens.

**Table S2 (separate file)**

**Noncoding element perturbation data**

(**A**) Data for E-G pairs generated by CRISPRi-FlowFISH screens in K562 cells.

(**B**) Data for E-G pairs generated by enhancer deletion followed by allele-specific RNA-seq in mESCs.

**Table S3 (separate file)**

**Dataset of experimentally tested noncoding element-gene connections**.

Including CRISPRi-FlowFISH screen and enhancer deletion data from this study as well as data from other studies perturbing noncoding elements with CRISPRi, 2-guide deletion, or other genome editing, and measuring the effect on gene expression through RNA-seq, allele-specific RNA-seq, CRISPR screens, or RT-qPCR.

(**A**) Enhnacer perturbation data in K562

(**B**) Enhancer perturbation data in other cell types.

**Table S4 (separate file)**

**Sequences of primers, gRNAs, siRNAs, and probesets**.

(**A**) Primer sequences for RT-qPCR and genotyping enhancer deletion clones

(**B**) gRNA sequences for single gRNA CRISPRi and enhancer deletion.

(**C**) Catalogue numbers (Thermo Fisher) for FlowFISH probesets.

(**D**) Catalogue numbers (Ambion) for siRNAs.

**Table S5 (separate file)**

**Sources of epigenomic and expression data**.

**Table S6 (separate file)**

**Genome-wide DE-G predictions computed using the ABC Model**

(**A**) Description of prediction file format and explanation of columns

(**B**) Predicted DE-G connections in K562 cells

(**C**) Predicted DE-G connections in GM12878 cells

(**D**) Predicted DE-G connections in primary erythroid progenitors

(**E**) Predicted DE-G connections in NCCIT cells,

(**F**) Predicted DE-G connections in LNCaP cells,

(**G**) Predicted DE-G connections in hepatocytes

(**H**) Predicted DE-G connections in mESC cells

(**I**) Genes predicted to have no distal enhancers

**Table S7 (separate file)**

**List of human ubiquitously expressed genes**

**Fig. S1.**
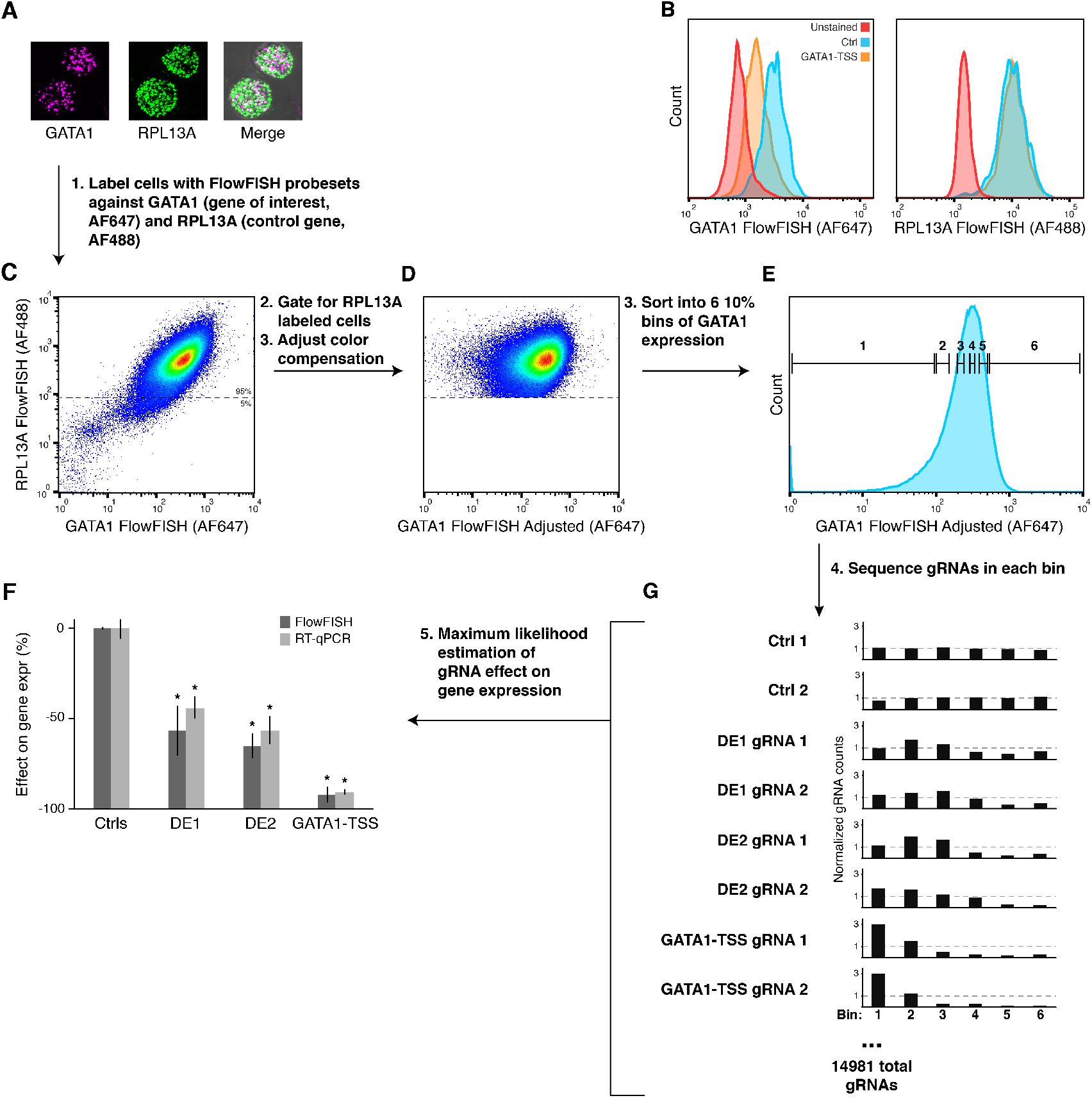
Sorting and sequencing strategy for CRISPRi-FlowFISH Screens. (**A**) K562 cells labeled with FlowFISH probesets against *RPL13A* (control gene) and *GATA1* (gene of interest) imaged by fluorescence microscopy. (**B**) Histograms of FlowFISH signal (arbitrary units of fluorescence) for *GATA1* (left) and *RPL13A* (right) in unlabeled K562s (red), K562s stained for *GATA1* expressing a gRNA against the *GATA1-TSS* (orange), or a non-targeting Ctrl gRNA (blue). (**C**) Scatterplot of FlowFISH fluorescent signal for *RPL13A* versus *GATA1*. (**D**) Cells in (C) with cells unstained for *RPL13A* (below dotted line in (C)) removed and using the color compensation tool to reduce the correlation between the control gene and gene of interest (see Methods). (**E**) Binning strategy for sorting FlowFISH-labeled cells into 6 bins each containing 10% of the cells. (**F**) Effect on gene expression as measured by CRISPRi-FlowFISH (dark grey) and RT-qPCR (light grey). Error bars: 95% confidence intervals for the mean of 2 gRNAs per target, 3505 Ctrl gRNAs for FlowFISH, and 6 Ctrl gRNAs for RT-qPCR, each measured in biological triplicate. *: *p* < 0.05 in t-test versus Ctrl. (**G**) Counts in each of the 6 bins for single gRNAs targeting the *GATA1* TSS, the two *GATA1* enhancers (DE1 and DE2) identified in Fulco *et al*. (*17*), and representative negative controls (Ctrl).

**Fig. S2.**
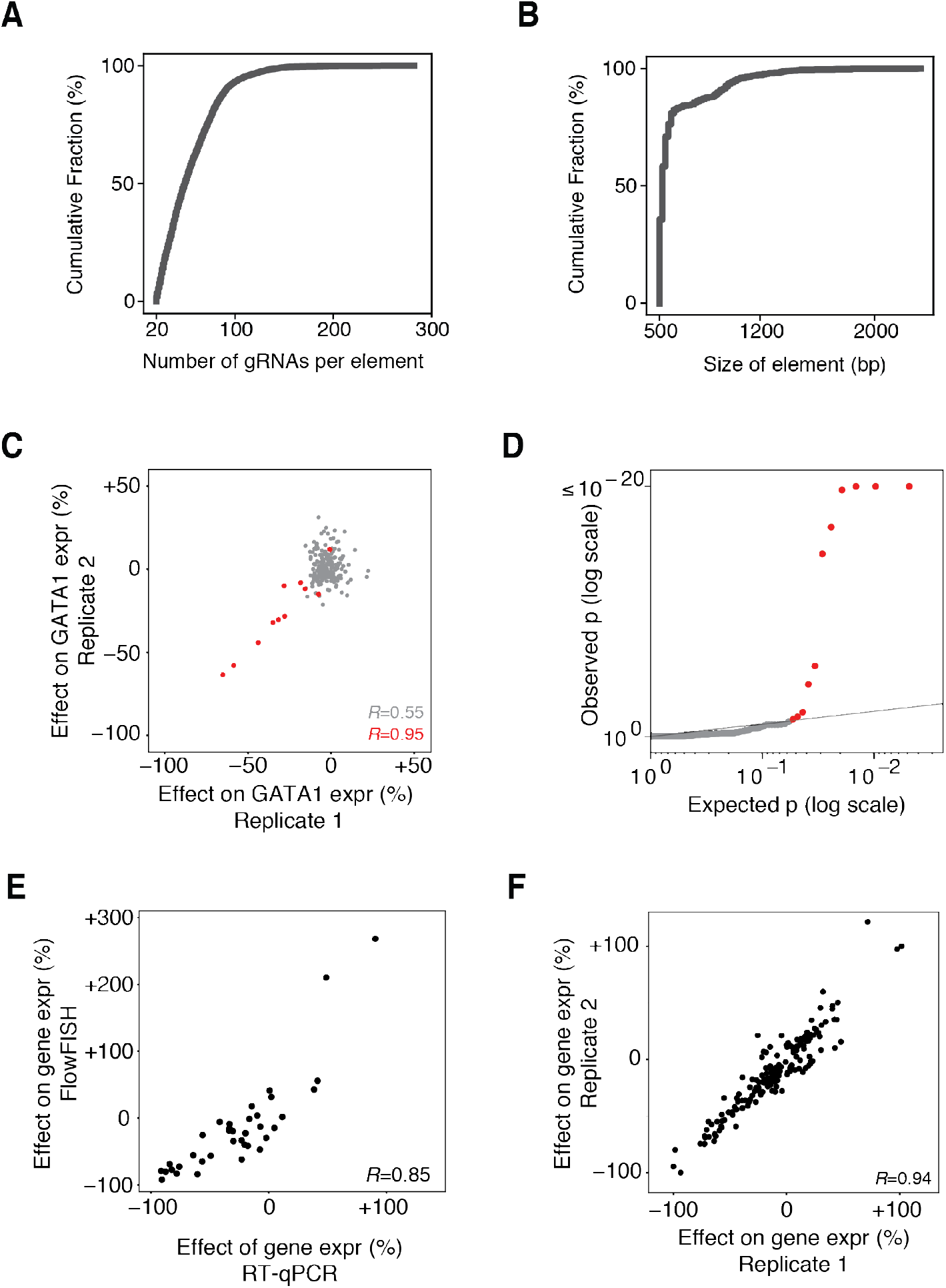
CRISPRi-FlowFISH reproducibly quantifies effects of regulatory elements. (**A**) Cumulative distribution plot of the number of gRNAs in each tested candidate element. (**B**) Cumulative distribution plot of the width of each tested candidate element. (**C**) Correlation between replicate CRISPRi-FlowFISH screens for *GATA1*. Red points denote elements significantly affecting expression. Pearson *R* = 0.95 for significant elements, 0.55 for all elements. (**D**) Quantile-quantile plot for *GATA1* CRISPRi-FlowFISH screen. Red points denote elements significantly affecting expression. Vertical axis capped at 10^−20^. (**E**) Correlation between effect on gene expression as measured by CRISPRi-FlowFISH screening and RT-qPCR for all 36 E-G pairs tested by both methods. Value is the mean effect of the two gRNAs for each element. (**F**) Correlation between effects on gene expression for all significant E-G pairs measured by replicate CRISPRi-FlowFISH screens.

**Fig. S3.**
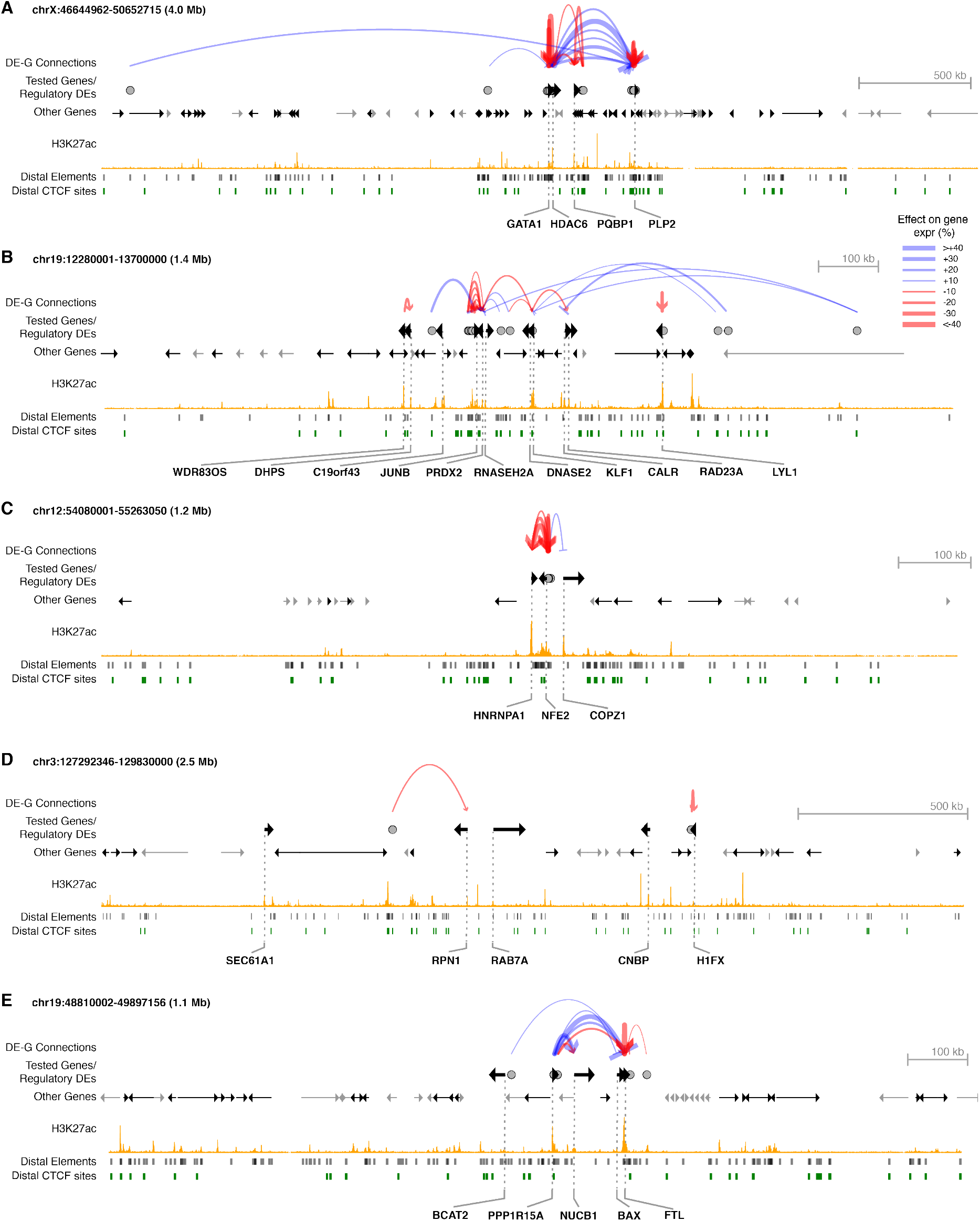
CRISPRi-FlowFISH enhancer perturbation dataset. DE-G connections are elements affecting the expression of the indicated gene in CRISPRi-FlowFISH screens in K562 cells. Red arcs denote activation, blue arcs denote repression. The width of the arc corresponds to the effect size. Distal elements (black) are tested DHS peaks. Distal CTCF elements (green) are CTCF ChIP-seq peaks within distal elements. Tested genes refer to genes for which we performed CRISPRi-FlowFISH experiments. Grey circles are DEs where perturbation with CRISPRi affects the expression of at least one tested gene as measured by CRISPRi-FlowFISH

**Fig. S4.**
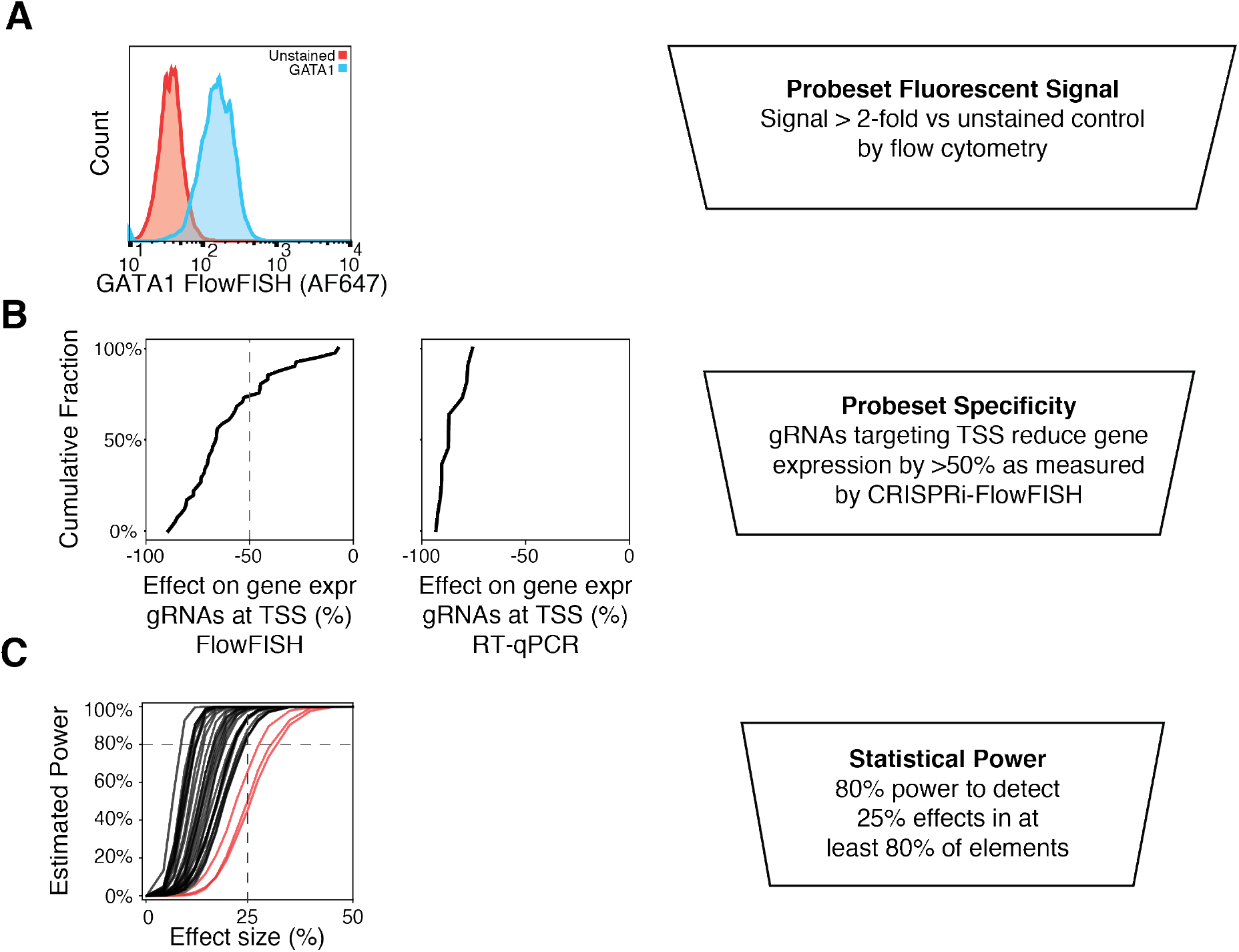
Quality filters for CRISPRi-FlowFISH probesets and screens. (**A**) Histogram of FlowFISH signal, as measured by flow cytometry, comparing K562 cells stained with *GATA1* probes compared to unstained, negative-control cells. We required probesets to have >2-fold mean fluorescent signal in stained versus unstained control. (**B**) Percent expression remaining in gRNAs targeting the TSS estimated from CRISPRi-FlowFISH screening. In all cases where we assessed CRISPRi knockdown by gRNAs at a TSS by qPCR, we observed >75% knockdown (right). However, some FlowFISH probesets reported <50% knockdown for gRNAs at their TSSs (left); we expect that some of the signal detected by these probesets results from off-target binding. Accordingly, we excluded these probesets from further analysis. (**C**) Power to detect a given effect size in 80% of E-G pairs for each gene. We analyzed those screens with at least 80% power to detect a 25% effect for at least 80% of tested elements. Red lines represent screens that did not meet this power threshold.

**Fig. S5.**
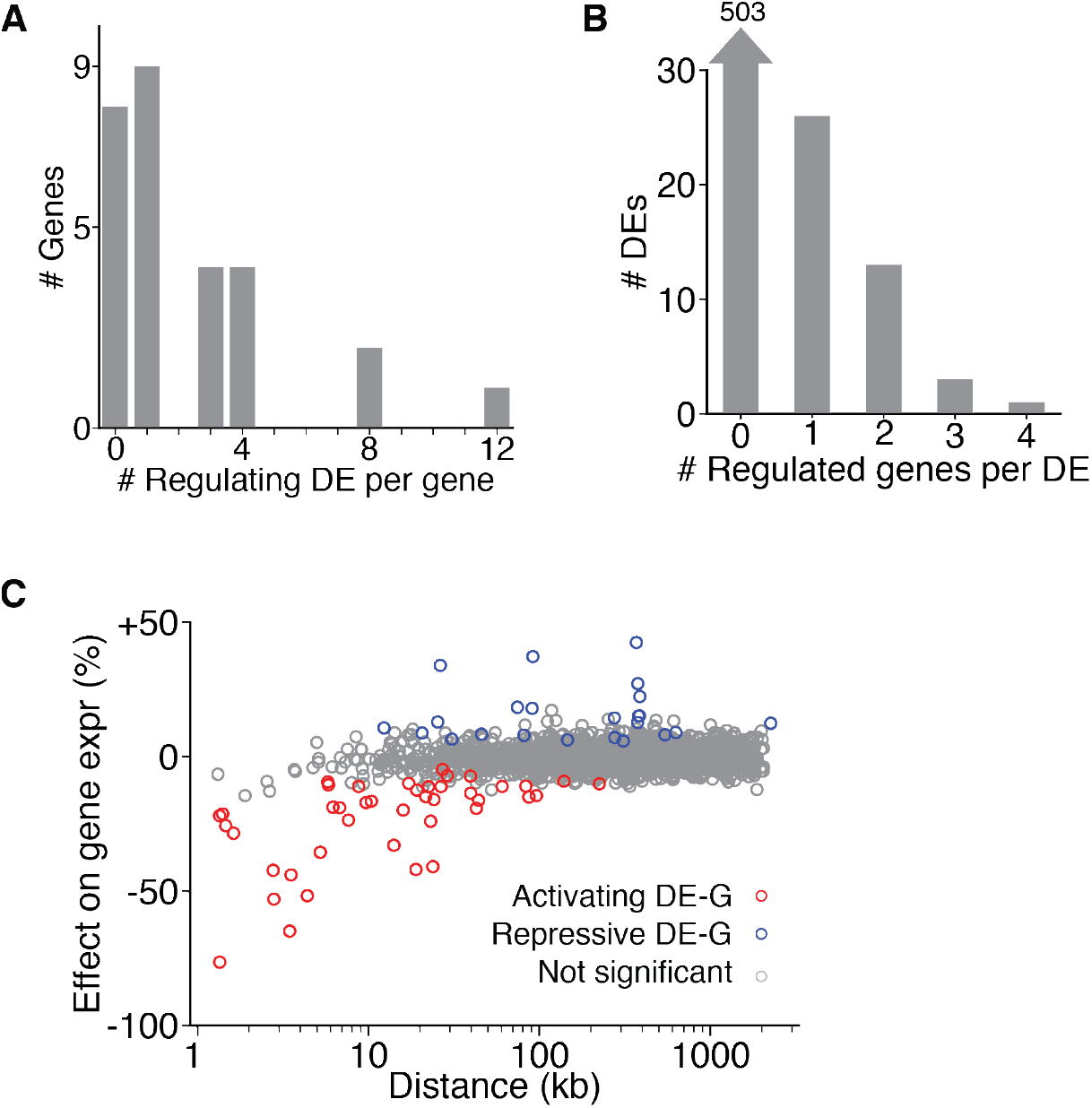
Properties of the CRISPRi-FlowFISH dataset. (**A**) Histogram of the number of distal elements affecting each gene in CRISPRi-FlowFISH experiments. (**B**) Histogram of the number of genes affected by each distal element tested in CRISPRi-FlowFISH experiments. (**C**) Comparison of genomic distance with observed changes in gene expression upon CRISPRi perturbation. Each dot represents one tested DE-G. Red dots: connections where perturbation resulted in a decrease in the expression of the tested gene. Blue dots: perturbation resulted in an increase. Grey dots: had no significant effect.

**Fig. S6.**
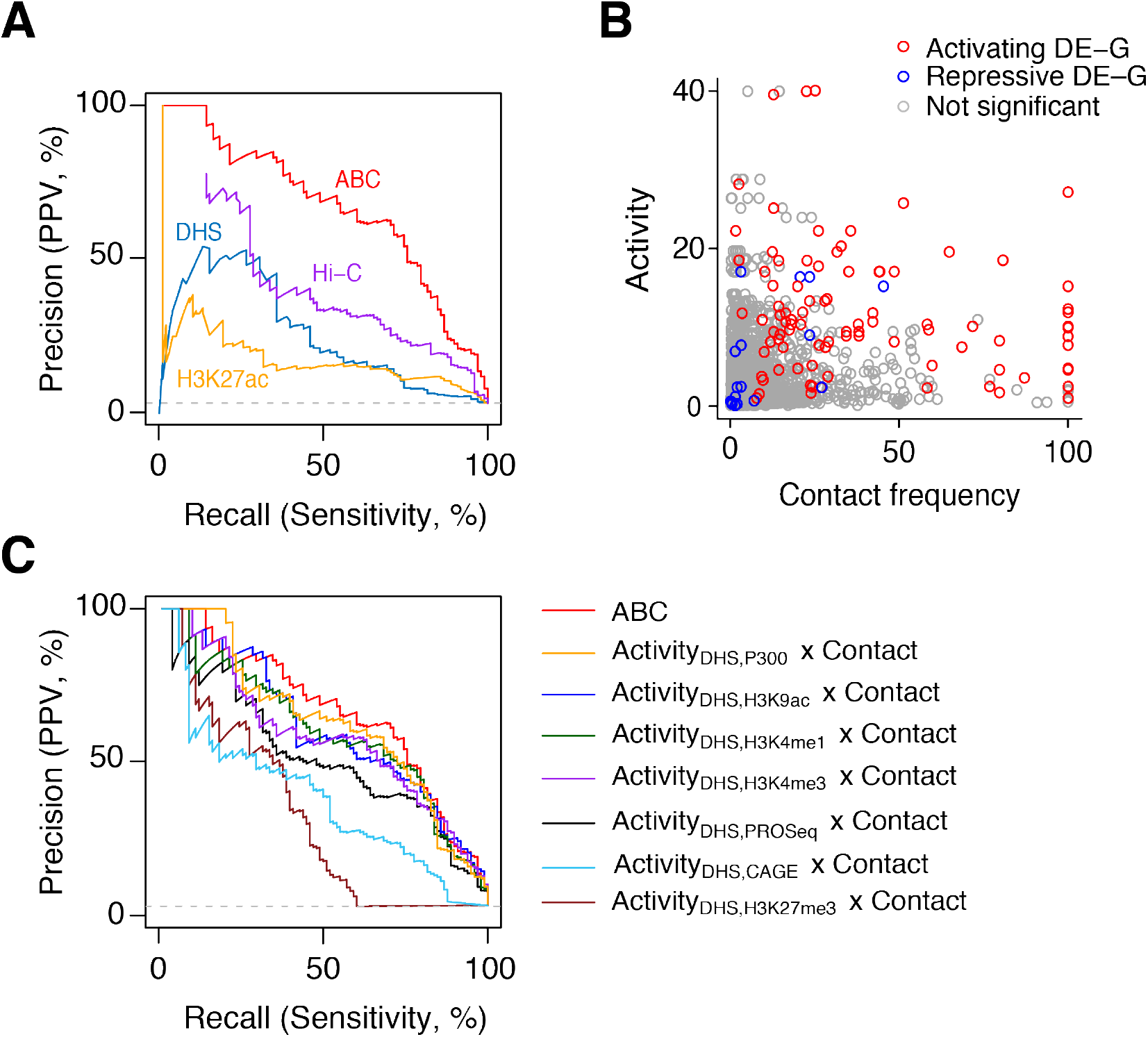
Comparison of ABC score to other predictors. (**A**) Precision-recall curves for classifying regulatory DE-G pairs, comparing each of the components of the ABC score. (**B**) Scatterplot of Activity and Contact frequency for each tested DE-G pair. KR-normalized Hi-C contact frequencies are scaled for each gene so that the maximum score of an off-diagonal bin is 100 (see Methods). (**C**) Precision-recall curves comparing different measures of Activity. Activity_Feature1,Feature2_ = sqrt(Feature1 RPM x Feature2 RPM). (ABC score corresponds to Activity_DHS,H3K27ac_ x Contact). Figure legends are ordered from best to worst AUPRC.

**Fig. S7.**
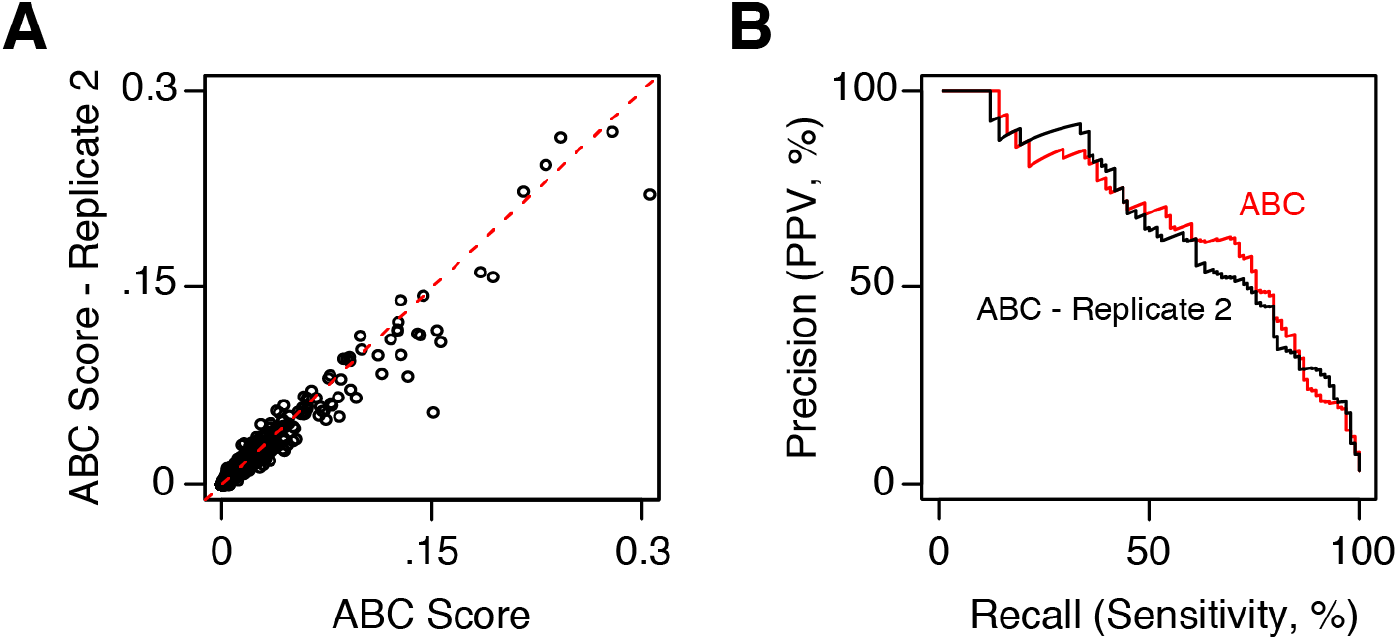
The ABC Score is reproducible between replicates of the epigenomic datasets. (**A**) Scatter plot of ABC Score computed using biological replicates for DHS and H3K27ac (Pearson *R* = .98). (**B**) Precision-recall curves for classifying regulatory DE-G pairs (Positive DE-G pairs are those where perturbation of element DE significantly reduces the expression of gene G) for the ABC Score using replicates 1 and 2 of DHS and H3K27ac.

**Fig. S8.**
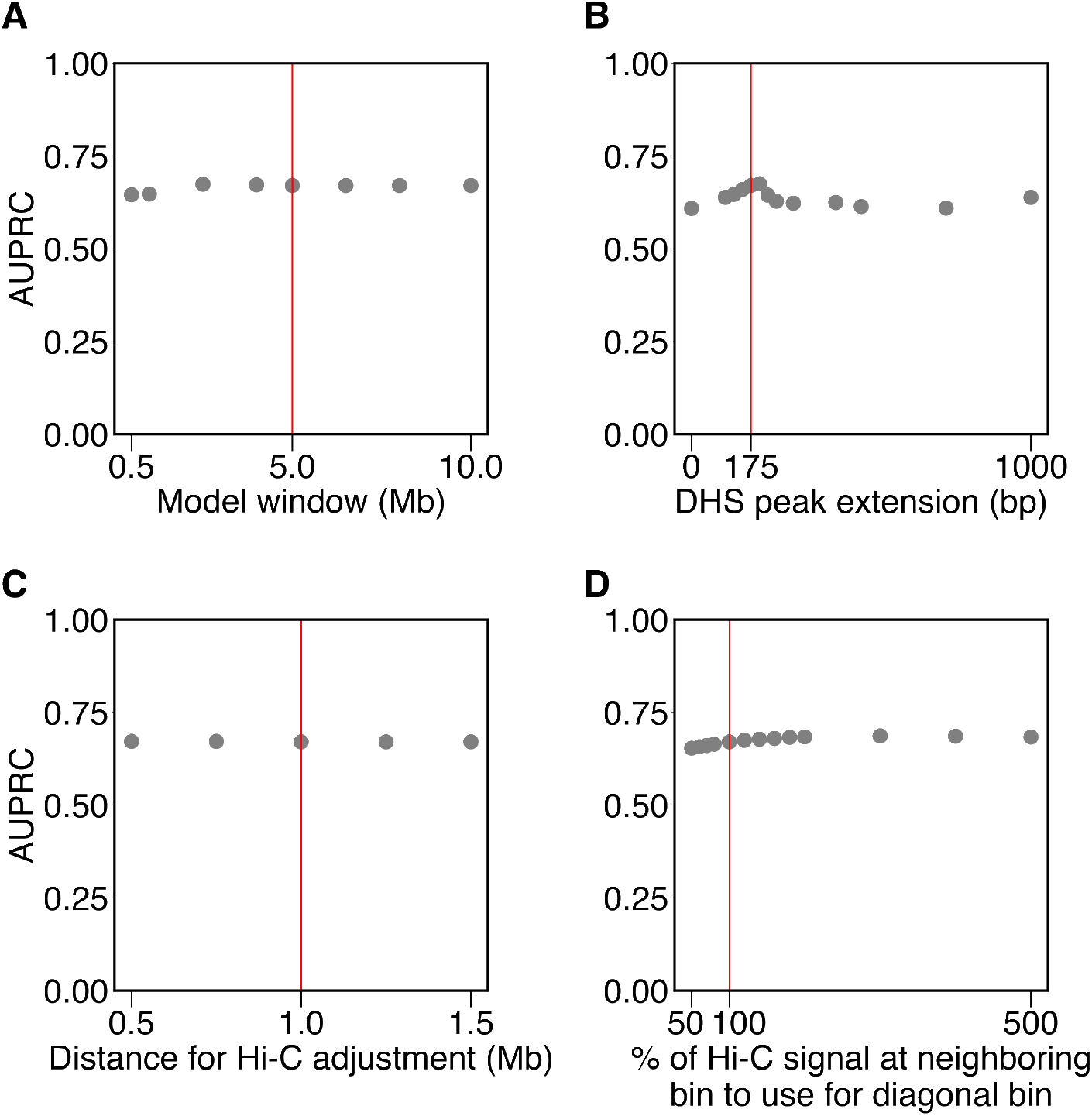
Sensitivity of ABC score performance to chosen parameters. Changing the parameters of the ABC score does not dramatically affect performance near the default values. Each panel presents the area under the precision recall curve (AUPRC) for the ABC score when changing the specified parameter. Red lines indicate the values used throughout this paper. (**A**) Genomic distance within which elements are included in the model. (**B**) Number of bases DHS peaks were extended on either side before merging to create candidate elements. (**C**) Genomic distance used to compute the pseudocount added to the Contact component (see Methods). (**D**) In processing of Hi-C data, each diagonal entry of the Hi-C matrix is replaced by the maximum of its four neighboring entries when estimating contact frequency at distances < 5 kb (see Methods).

**Fig. S9.**
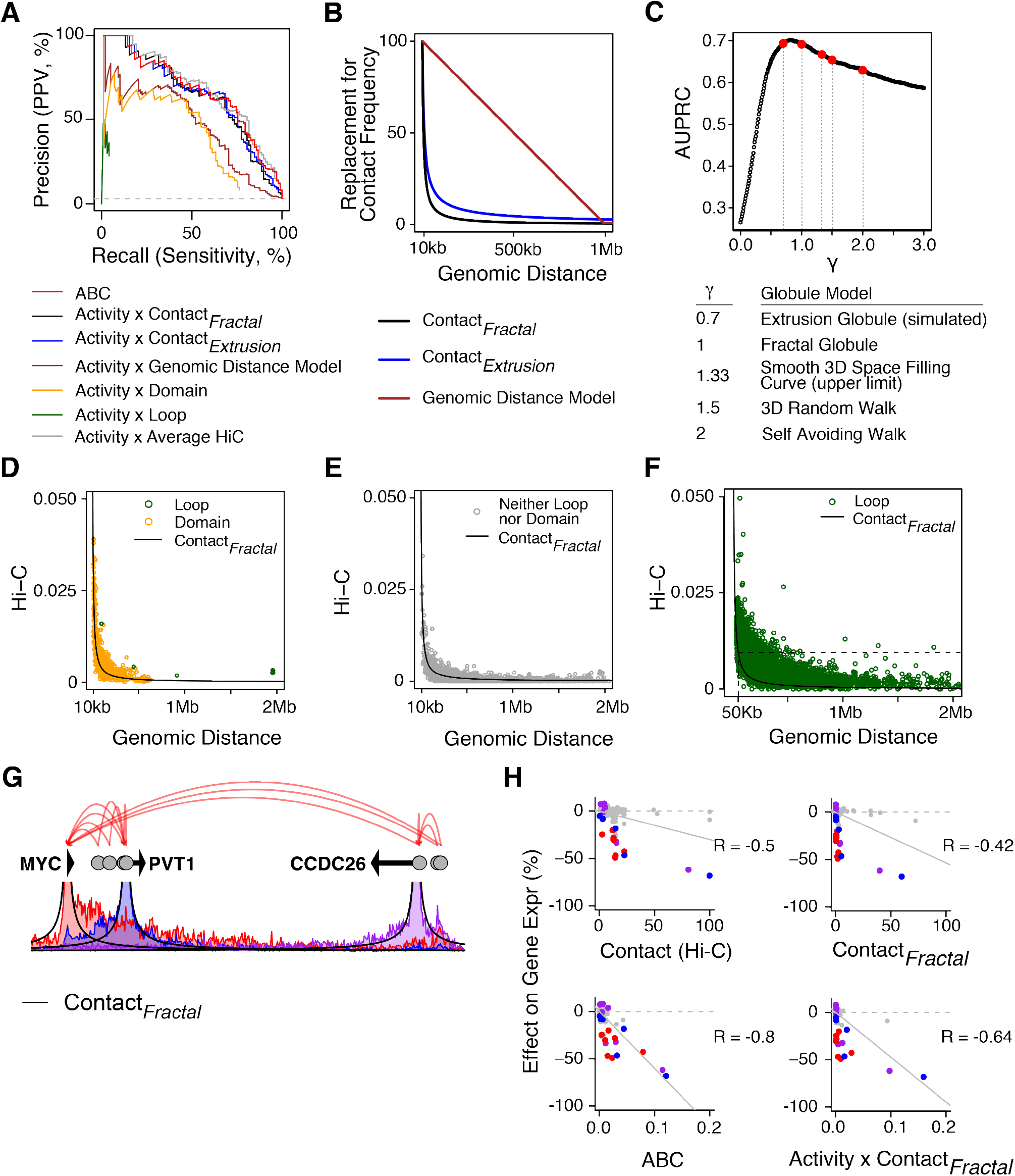
Testing other methods to estimate contact frequency for the ABC score. (**A**) Precision-recall curves comparing the ABC score to other models where the Contact component is replaced with binary Hi-C features (loops or domains) or decreasing functions of genomic distance (as visualized in panel B). *Activity x Genomic Distance:* Contact component is proportional to max(.01, (1e6 - *Distance*)/1e6). *Activity x Contact_Fractal_, Activity x Contact_Extrusion_*: Contact component is proportional to *Distance*^−γ^. *Contact_Fractal_* uses γ = 1, *Contact_Extrusion_* uses γ = 0.7. *Activity x Loop* and *Activity x Domain:* Contact component replaced by 1 if the element and gene TSS are located at the anchors of the same loop or within the same contact domain, respectively, or 0 otherwise. (**B**) Visualization of the quantitative functions used in (**A**) to replace contact frequency. Y-axis is in arbitrary units. In models of chromosome dynamics that assume chromatin is a featureless, uniform polymer in the globular state, Contact is inversely proportional to genomic distance raised to a fixed power (γ). Extrusion globule and fractal globule models (γ = 0.7 and 1) well represent the empirically observed Hi-C contacts at various distances (*38*). (**C**) AUPRC for ABC models where the Contact component is replaced with *Distance*^−γ^, with γ in the range [0, 3]. Values of γ corresponding to various polymer models are highlighted in red. The optimal values of γ as estimated from our CRISPRi data correspond to the values of γ that best predict Hi-C data (in the range of 0.7-1) (*38*). (**D, E**) Scatterplot of genomic distance vs contact frequency (Hi-C) for K562 tested DE-G pairs whose distance is greater than 10 kb. Colors represent membership in the same contact domain (orange), Hi-C loop (green) or neither annotation (gray). These relationships explain why the ABC score performs similarly to the Activity x *Contact_Fractal_* model: the power law relationship explains 69% of the variance of Hi-C contact frequency. In contrast, the ABC score performs very differently from the Activity x Loop and Activity x Domain models because loops and domains are not predictive of contact frequency. Y-axis is KR-normalized Hi-C signal (and, for convenience, is not scaled on a per-gene basis as is used in ABC model, see Methods). (**F**) Scatterplot of genomic distance vs quantitative contact frequency (Hi-C) for all loops in K562 (*59*). Although Hi-C contact frequency at loops is higher than expected under the Fractal Globule model, the absolute increase in contacts is modest. For example, the loops with highest contact frequency at 500 kb have the expected contact frequency of non-loop loci at 50 kb (dotted line). (**G, H**) Comparison of DE-G predictions in the MYC locus using Hi-C vs the *Contact_Fractal_* model (G) Visualization of Hi-C tracks anchored at the MYC, PVT1 and CCDC26 promoters (colored lines), compared to the *Contact_Fractal_* model (black lines). (H) Computation of the ABC score for DE-G pairs in the MYC locus using Hi-C vs the *Contact_Fractal_* model. Using Hi-C data better predicts the quantitative effects of enhancers in this locus.

**Fig. S10.**
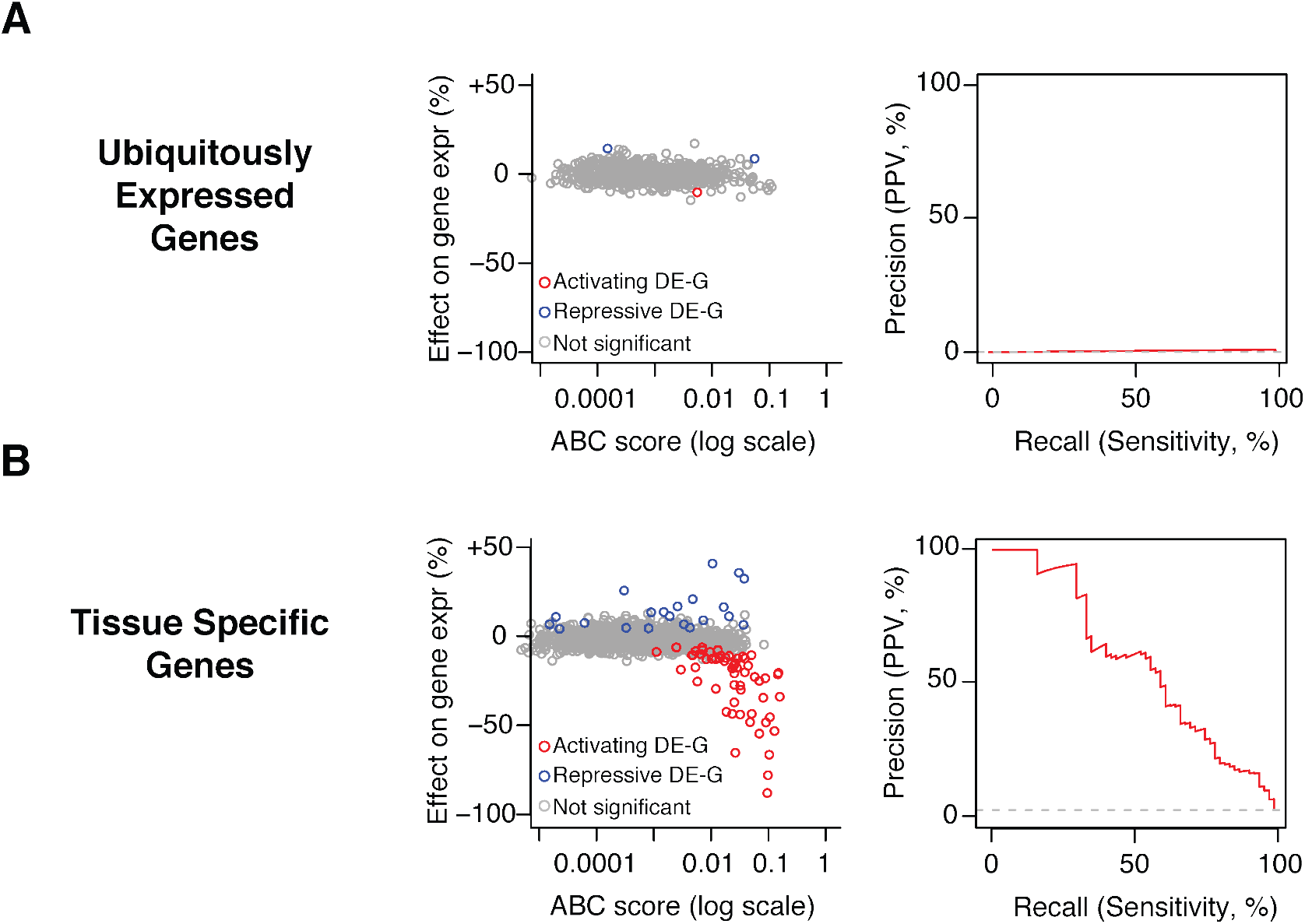
Tissue-specific genes have more distal enhancers than ubiquitously-expressed genes. (**A**) Left: Comparison of ABC scores (predicted effect) with observed changes in gene expression upon CRISPR perturbations. Each dot represents one tested DE-G pair where G is a ubiquitously-expressed gene. Right: precision-recall curve for ABC score in classifying regulatory DE-G pairs where each G is a ubiquitously-expressed gene. (**B**) Same as (**A**) for tissue-specific genes. All panels include only the subset of our dataset for which we have CRISPRi tiling data to comprehensively identify all enhancers that regulate each gene (28 genes from this study, 2 from previous studies).

**Fig. S11.**
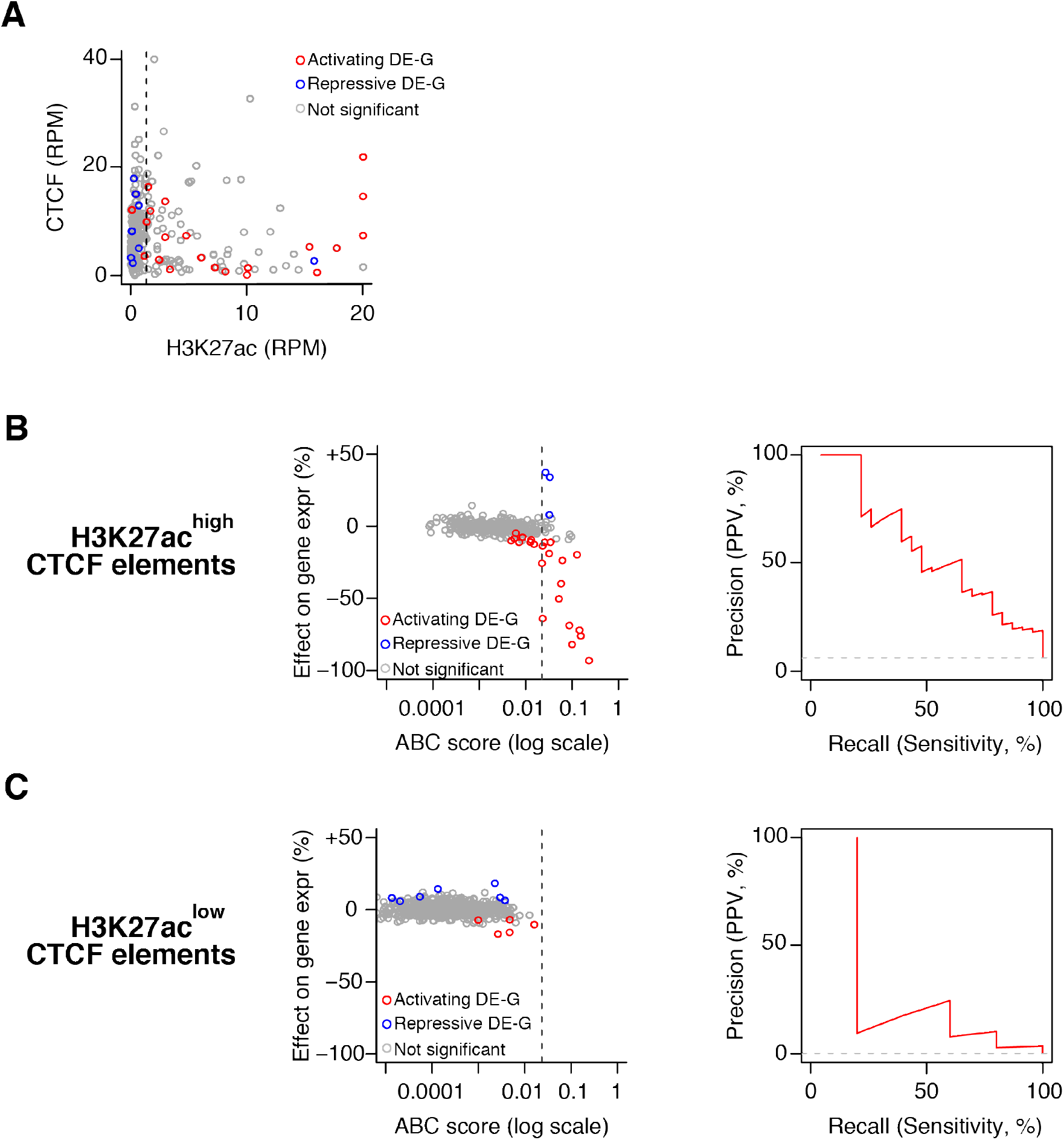
Analysis of CTCF-bound elements. (**A**) Scatterplot of CTCF signal (reads per million) vs H3K27ac signal (reads per million) for all DE-G pairs where the DE is bound by CTCF (see Methods). Dotted black line corresponds to the median H3K27ac signal for all distal elements in the dataset. We denote elements whose H3K27ac signal is greater than the median “H3K27ac^High^ CTCF elements” and those with H3K27ac signal less than the median “H3K27ac^Low^ CTCF elements”. (**B**) Left: comparison of ABC scores (predicted effect) with observed changes in gene expression upon CRISPR perturbations. Each dot represents one tested DE-G pair where the DE is a H3K27ac^High^ CTCF element. Right: precision-recall curve for the ABC score in classifying regulatory DE-G pairs where each DE is a H3K27ac^High^ CTCF element. (**C**): Same as (B) for H3K27ac^Low^ CTCF elements.

**Fig. S12.**
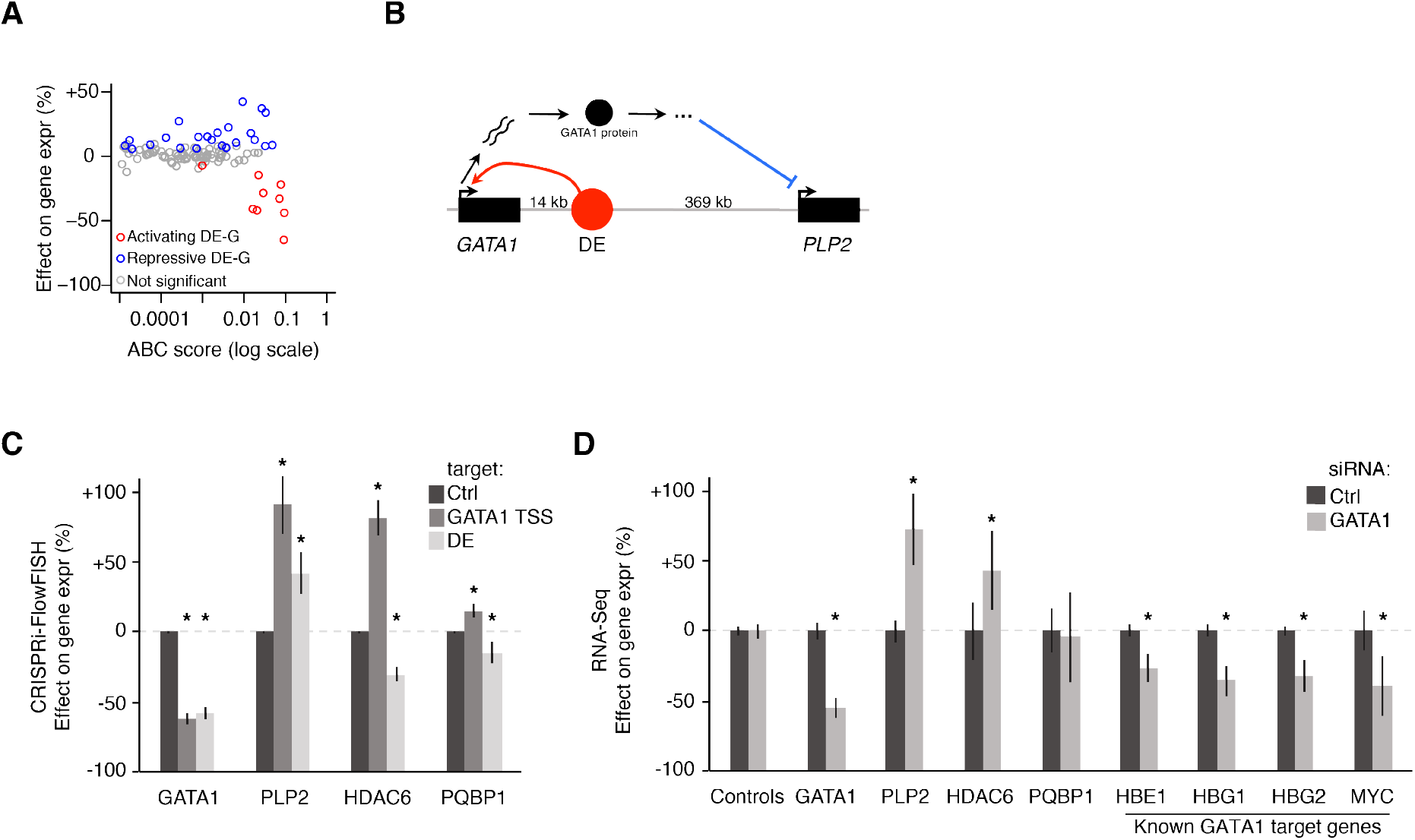
Elements that repress a distal gene are likely explained by indirect regulatory effects. (**A**) Comparison of ABC scores (predicted effect) with observed changes in gene expression upon CRISPR perturbations. Each dot represents one tested DE-G pair where the element represses at least one gene. (**B**) Summary of the effect of a *GATA1*-regulating DE on *PLP2*. The observed repressive effect of this DE on *PLP2* is consistent with this DE activating *GATA1* (red arc), which in turn represses *PLP2* via a transacting function of the GATA1 protein product (blue arc). (**C**) Effects of inhibiting *GATA1* TSS or a *GATA1* enhancer (DE) with CRISPRi. mRNA expression measured by CRISPRi-FlowFISH. Error bars: 95% confidence intervals for the mean of all gRNAs within the target element (Table S3). *: BH-adjusted *p* < 0.05 in t-test versus negative controls (see Methods). (**D**) Effects of inhibiting *GATA1* with siRNAs on gene expression of *GATA1, PLP2, HDAC6, PQBP1*, and known GATA1 transcription factor targets (*8, 79, 80*) as measured by RNA sequencing of cells transfected with *GATA1* siRNA compared to nontargeting siRNAs (Ctrl). Control genes are the average of commonly used housekeeping genes (*ACTB, B2M, C1orf43, CHMP2A, EMC7, GAPDH, GPI, PGK1, PPIB, PSMB2, PSMB4, REEP5, RPL13A, SNRPD3, TBP, TUBB, VCP*, and *VPS29*). Error bars: 95% confidence interval for the mean of two siRNAs with four independent transfections each. *: BH-adjusted *p* < 0.05 from DESeq2 for *GATA1* siRNA versus Ctrl (see Methods).

**Fig. S13.**
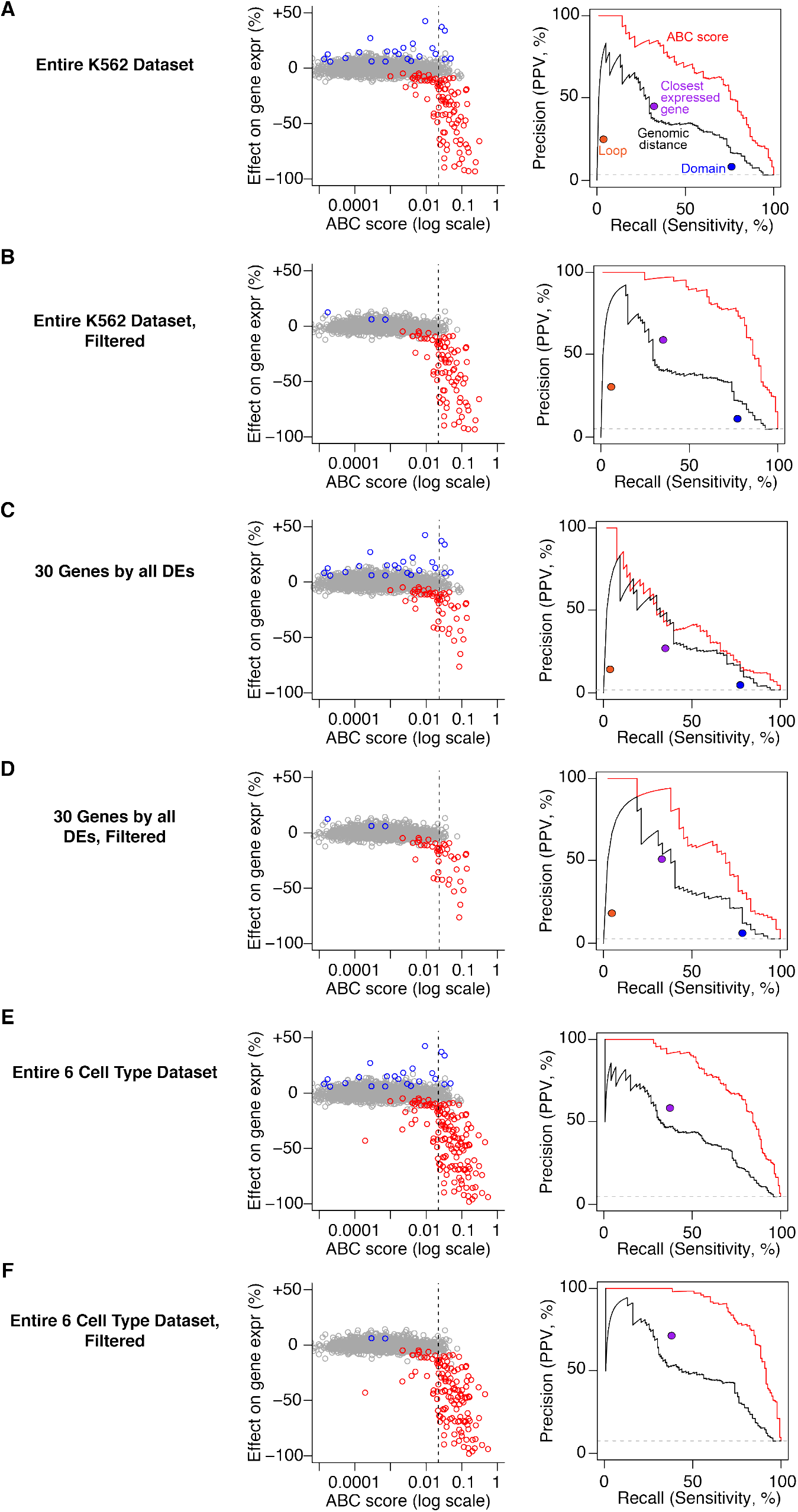
Performance of the ABC score after filtering ubiquitously expressed genes, CTCF elements, and indirect effects. Performance of the ABC score on subsets of the CRISPR dataset. (**A**) Entire initial dataset in K562 cells (same as Fig 3). (**B**) K562 dataset with H3K27ac^low^ CTCF elements, DE-G pairs likely to result from indirect effects, and ubiquitously expressed genes removed. (**C**) DE-G pairs in CRISPRi tiling experiments that, for a given gene, perturb and test the effects of all nearby DEs. (**D**) Subset described in (C) with H3K27ac^low^ CTCF elements, DE-G pairs likely to result from indirect effects, and ubiquitously expressed genes removed. (**E**) Entire dataset across 6 cell types. Includes cell types without Hi-C data, so the performance of Hi-C loops and domains cannot be evaluated. (**F**) Subset described in (E) with H3K27ac^low^ CTCF elements, DE-G pairs likely to result from indirect effects, and ubiquitously expressed genes removed. In each panel: Left plot is a comparison of ABC scores (predicted effect) with observed changes in gene expression upon CRISPR perturbations. Each dot represents one tested DE-G pair. Right plot is a set of precision-recall curves for classifying regulatory DE-G pairs (Positive DE-G pairs are those where perturbation of element DE significantly reduces the expression of gene G).

**Fig. S14.**
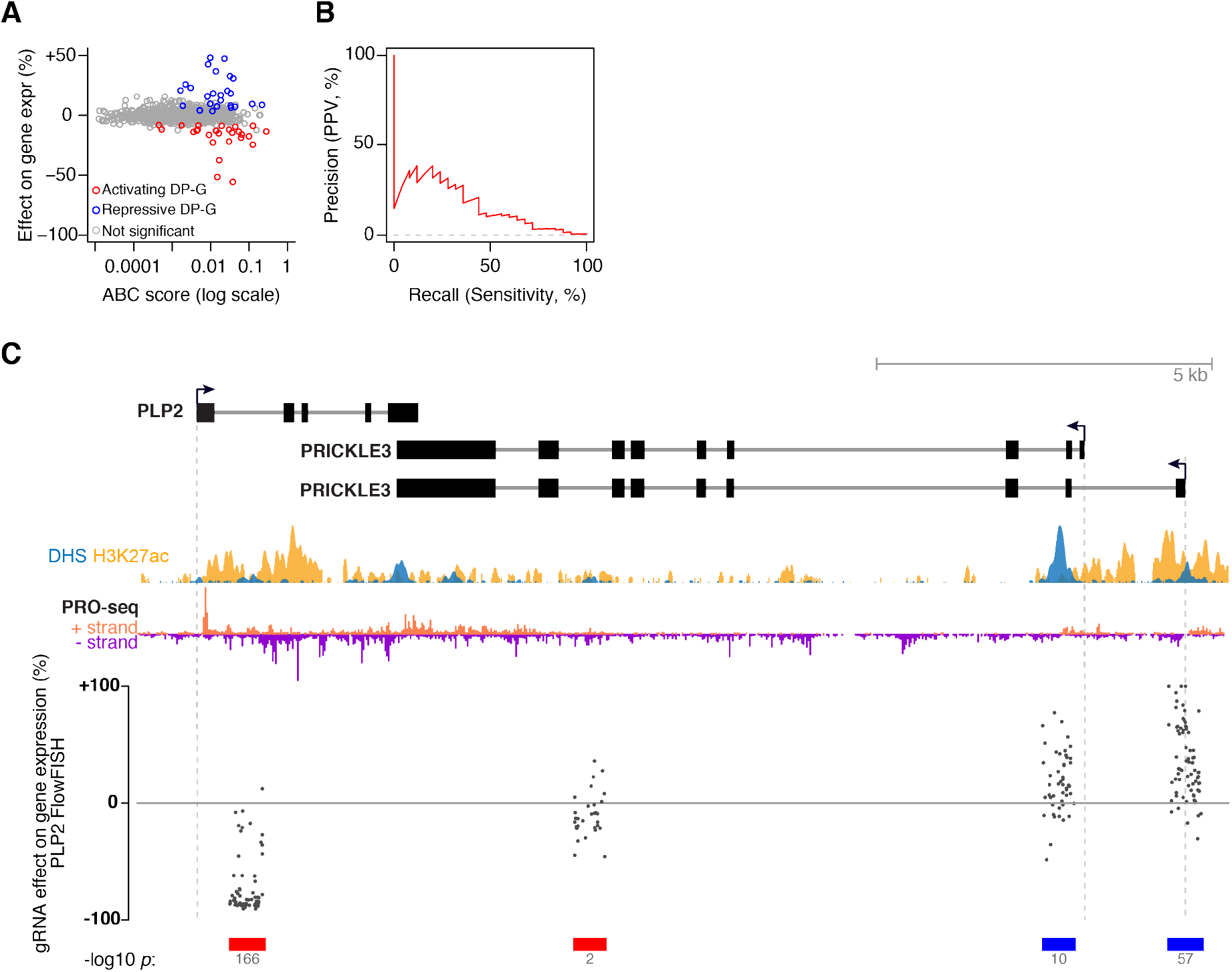
Effects of promoters on nearby genes. (**A**) Comparison of ABC scores (predicted effect) with observed changes in gene expression upon CRISPR perturbations in K562 cells. Each dot represents one tested DP-G pair (where the element itself is a promoter). (**B**) Precision-recall curve for classifying regulatory DP-G pairs (Positive DP-G pairs are those where perturbation of promoter P significantly reduces the expression of distal gene G). (**C**) Some promoters appear to affect expression of neighboring genes by transcriptional interference. One example is the effect of *PRICKLE3* on *PLP2*. Points represent the effect of gRNAs on *PLP2* expression, as measured by CRISPRi-FlowFISH. Vertical axis capped at +100%. Red and blue bars: DHS elements in which CRISPRi leads to a significant decrease (red) or increase (blue) in *PLP2* expression. Transcription of *PRICKLE3* as measured by PRO-seq (negative strand, purple) extends into the gene body of *PLP2* (positive strand, salmon). Therefore, transcriptional interference may explain why CRISPRi inhibition of the *PRICKLE3* promoter leads to an increase in *PLP2* expression.

**Fig. S15.**
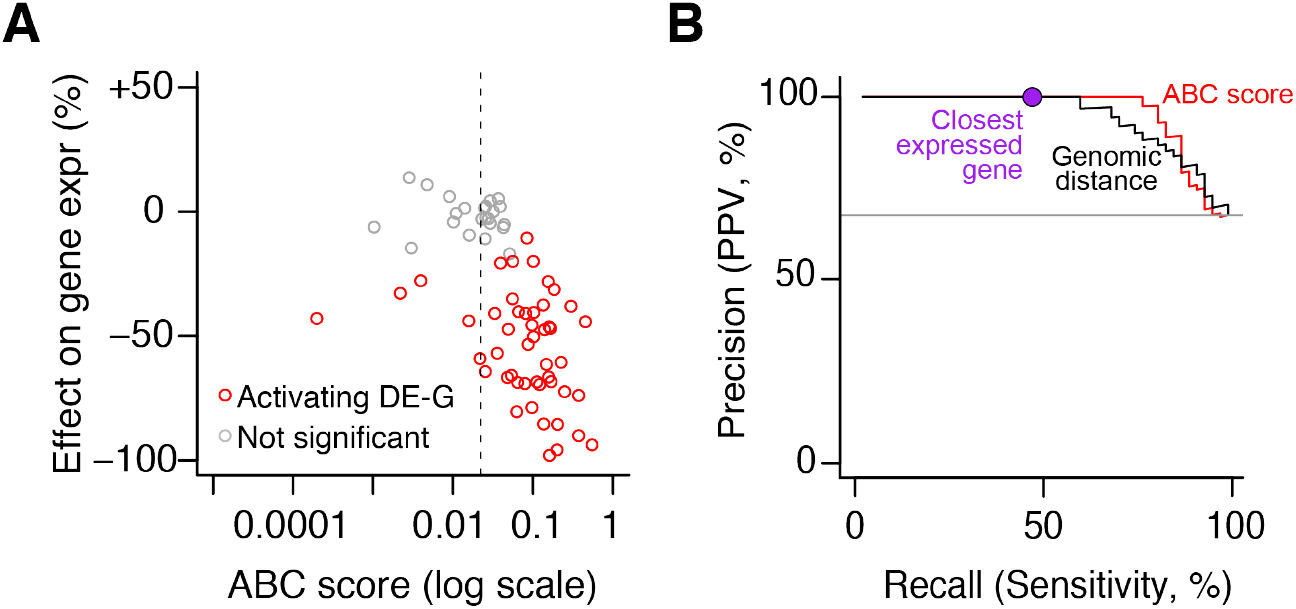
The ABC model generalizes across cell types. (**A**) Comparison of ABC scores (predicted effect) with observed changes in gene expression upon perturbations in GM12878, LNCaP cells, NCCIT cells, primary human hepatocytes, and mouse ES cells. Each dot represents one tested DE-G pair. (**B**) Precision-recall plot for classifiers of DE-G pairs shown in (A). Positive DE-G pairs are those where the distal element significantly decreases expression of the gene. Curves represent the performance for predicting significant decreases in expression for DE-G pairs based on thresholds on the ABC score (red) and genomic distance between the DE and the TSS of the gene (black). The purple circle represents the performance of assigning each DE to the closest expressed gene. DE-G pairs that were not significant are filtered for those that pass the same stringent power filter applied to the K562 dataset, requiring 80% power to detect a 25% effect on gene expression. (See Methods. See Fig 4 for data in these types using a lenient power filter of 80% power to detect a 50% effect on gene expression).

